# Urbanization weakens the latitudinal diversity gradient in birds

**DOI:** 10.1101/2025.08.04.668553

**Authors:** Jory Griffith, Jennifer M. Sunday, Anna L. Hargreaves

**Affiliations:** Department of Biology, McGill University; Montreal, Canada

## Abstract

The latitudinal diversity gradient of increasing species richness toward the tropics is Earth’s most ubiquitous and ancient macroecological pattern, but may not be immune to modern anthropogenic change. Urbanization represents a particularly severe and globally-distributed ecological change; while urbanization generally reduces local diversity, whether it affects large-scale diversity patterns is unclear. Analyzing global databases including >10,000 bird species on five continents, we show that the avian latitudinal diversity gradient is 3x weaker in urban than natural areas. Urbanization reduces diversity across latitudes, but tropical cities lose disproportionately more species, partly because tropical bird assemblages contain more specialists and specialists are particularly excluded by urbanization. Our results show that urbanization is disrupting macroecological patterns, and suggest that specialization mediates both latitudinal diversity gradients and urbanization impacts.

## Main Text

The dramatic increase in biodiversity from the poles to equator is one of the world’s oldest and most striking biogeographic patterns (*1*). This latitudinal diversity gradient is pervasive across continents, realms, and taxonomic groups, and has inspired fundamental theory in ecology and evolution about the processes governing diversity (*2–4*). However, despite being extensive and ancient, the latitudinal diversity gradient may not be immune to the effects of anthropogenic change, as human activity affects species diversity around the globe (*5–7*).

A particularly widespread and intensive form of anthropogenic change is urbanization. Although cities cover only 0.7% of global land area (*8*), they contain more than half the human population and are growing rapidly (*9*). Globally, the habitat loss, degradation, and fragmentation caused by urbanization have reduced species richness in cities by as much as 60% compared to nearby natural areas, although effects vary among taxa (*10–12*). However, while local effects of urbanization have been extensively studied, we have only just begun to test how these scale up to influence macroecological patterns as fundamental as the latitudinal diversity gradient (*13–15*).

Given that urbanization generally reduces diversity (*11, 16*), we explore three alternative hypotheses for how urbanization could affect the latitudinal diversity gradient (Fig. 1A). Urbanization could cause a similar per-area species loss across latitudes (‘*uniform numeric loss*’ hypothesis). In this scenario, urbanization would reduce diversity uniformly across latitudes but would not affect the strength (i.e. slope) of the latitudinal diversity gradient. While a uniform reduction in diversity may be intuitive (*13*), there is not a clear ecological mechanism by which a similar number of species would be lost from areas with very different species diversities.

**Fig. 1.**
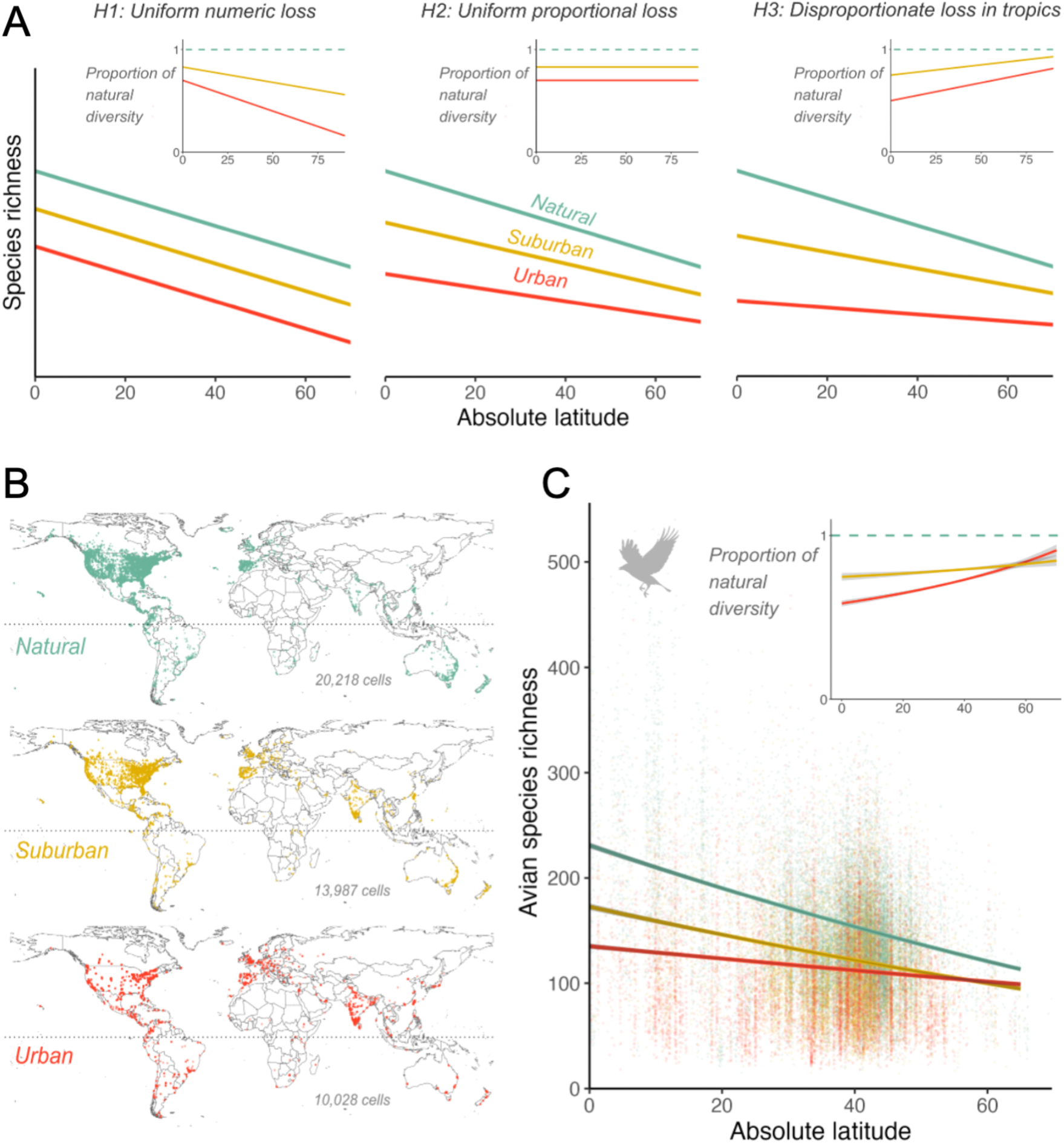
Urbanization weakens the latitudinal diversity gradient. **A**) Hypotheses for how urbanization might affect latitudinal diversity gradients, by excluding a uniform number (*H1*), uniform proportion (*H2*), or disproportionate number (*H3*) of species at low vs. high latitudes. Insets show diversity in urban and suburban areas as a proportion of the diversity in natural areas, to clearly differentiate *H2* and *H3*. **B**) Global distribution of data in main analyses.

Alternatively, urbanization might exclude a similar proportion of species across latitudes (‘*uniform proportional loss*’ hypothesis), for example if urbanization excludes species with certain habitat requirements (*17*), and the proportion of species with these requirements is relatively constant across latitude. In this scenario, urbanization would weaken the latitudinal diversity gradient by filtering out more species from species-rich communities at low-latitudes.

Finally, lower latitudes might contain a greater proportion of species who cannot coexist with urbanization, such that urbanization excludes a higher proportion of species toward the tropics (‘*disproportionate loss from tropics*’ hypothesis). For example, urbanization tends to exclude species with more specialized habitat and food requirements (*18, 19*), and lower-latitude communities are hypothesized to contain a higher proportion of specialized species (*20–22*). The disproportionate loss scenario could severely reduce or even eliminate the latitudinal diversity gradient in urban areas, meaning urbanization impacts not only local richness, but the biogeographic patterns that have defined the distribution of global biodiversity for millennia.

To test among these hypotheses, we compared latitudinal patterns of bird diversity among natural, suburban, and urban areas around the world. Bird diversity shows a strong latitudinal gradient (*23*) and birds have the wealth of fine-scale occurrence data needed to compare global diversity patterns. Urbanization often reduces local avian richness (*16, 24, 25*) and seems to favor species that have less specialized diet, habitat, or foraging niches (*19*). Previous attempts to examine how urbanization affects avian latitudinal diversity gradients have been limited to relatively small spatial and temporal scales and have yielded conflicting results (*26, 27*), underscoring the need for global, multi-year data to provide strong tests of latitudinal patterns (*28*).

We also explored the role of seasonality on global avian diversity patterns. Roughly 20% of bird species winter at low latitudes but migrate to higher latitudes to breed. Migration is especially common in the northern hemisphere, thus the latitudinal diversity gradient may vary seasonally within each hemisphere (*29, 30*), being strongest in winter and weakest in summer. In contrast, the negative effects of urbanization on diversity may be strongest in summer, as during winter many temperate bird species are more urban tolerant (*18*), and may even prefer urbanized habitats for their greater warmth or food availability (*31, 32*). Combined, these effects should particularly weaken the latitudinal diversity gradient in summer urban areas.

To test whether urbanization is disrupting the latitudinal diversity gradient in birds, we synthesized global data across 143 countries and 10,163 bird species (Fig. 1B). We characterized bird species richness within 1×1 km grid cells using species checklists from eBird, a database of citizen science observations with global coverage and the metadata needed to standardize for observation effort. We used databases on human land use to categorize each 1×1 km cell as natural, suburban, or urban, then tested whether bird diversity and the latitudinal diversity gradient differed among these categories. We explored whether results differed among seasons or hemispheres, since both could affect diversity patterns. Finally, using available estimates of species’ habitat and diet breadth, we tested whether urbanization disproportionately excludes specialized species (i.e. those with narrower habitat or diet breadth), and whether this results in disproportionate loss of avian diversity from cities at lower latitudes.

## Results and Discussion

We found that urbanization strongly reduces both overall diversity and the latitudinal diversity gradient in birds (Fig. 1C, Table S1). Globally, urban areas had 32% fewer bird species than natural areas (mean 95 vs. 141 species/km^2^), with suburban richness intermediate (mean 114 species/km^2^). Consistent with the “*disproportionate loss from tropics*” hypothesis, urbanized areas were missing a higher proportion of species at tropical than temperate latitudes (Fig. 1C). As a result, the overall latitudinal diversity gradient was 3x steeper in natural than urban areas (a gain in mean diversity of 118 vs. 36 species from 65° to 0°, respectively; Table S2). Indeed, from 65° to the equator, mean diversity more than doubled in natural areas (increase of 104%) but increased by only 36% in urban areas (Table S2). The disproportionate loss hypothesis was supported in both the northern and southern hemispheres (Fig. S1), and if we analyzed diversity in 5 km cells (Fig. S2), used different thresholds to define ‘natural’ cells (Fig. S3), or used different minimum sampling efforts for cell inclusion (Fig. S4). Thus urbanization has not only reduced local bird diversity across latitudes, but has weakened one the world’s more prominent macroecological gradients.

Coloured points are 1×1 km grid cells that could be classified as natural, suburban, or urban and had adequate bird occurrence data to estimate diversity. **C**) Observed avian diversity. Bird diversity increases toward the tropics in natural (top line), suburban, and urban (bottom line) areas, but the diversity gradient was significantly weaker in urbanized areas (statistical results in Table S1). Inset: urban and suburban areas lost a greater proportion of species at lower latitudes, supporting *H3*. Lines, grey polygons, and points show means, 95% confidence intervals, and partial residuals (narrow CI sometimes obscured by the trend line; see Table S9 for values). Values are marginal means, i.e. averaged across other model predictors, including hemisphere.

Urbanization’s dampening effect on the latitudinal diversity gradient was at least partly driven by greater loss of specialist species toward lower latitudes. On average, species had narrower habitat and diet niches in lower compared to higher latitudes (Fig 2; Table S3), providing some of the largest-scale support to date for the long-standing hypothesis that species are more specialized in the tropics. Within each latitudinal zone, bird species who were never found in urban areas had more specialized habitats (Fig. 2) and diets (Fig. S5) than species who could be found in urban areas, suggesting that cities disproportionately exclude specialists. Finally, the difference in specialization between urban and non-urban birds was highest in the tropics, and decreased toward the poles. All of these results held across seasons (Fig. S6; Table S4). Together, this evidence suggests a mechanism for urbanization’s disproportionate effect on bird diversity in the tropics; tropical latitudes contain more specialized species, and these specialists are disproportionately excluded from urban environments.

**Fig. 2.**
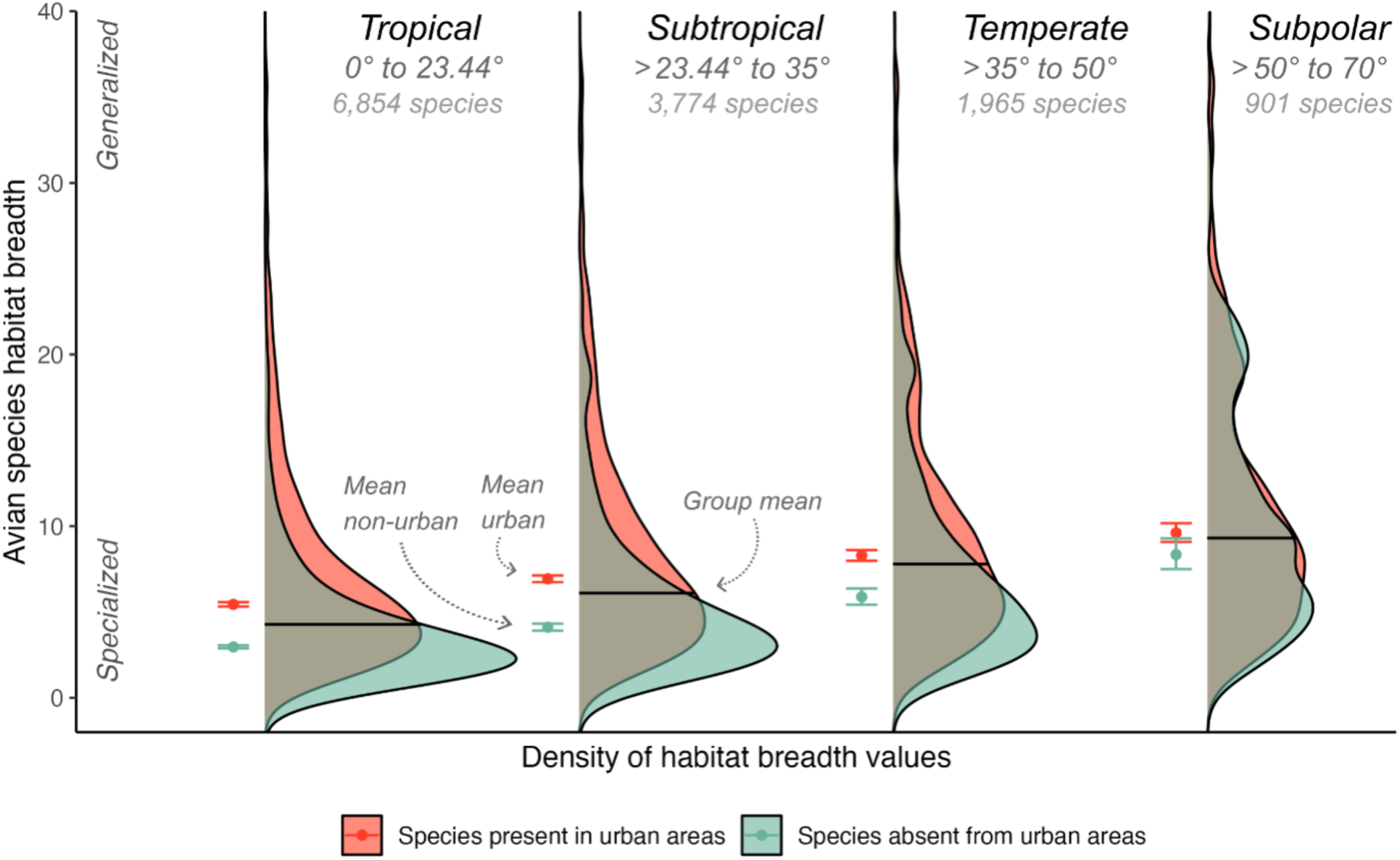
Specialized birds are disproportionately missing from urban areas, especially in the tropics. Distribution of habitat-breadth scores in each of four latitudinal zones for bird species found in urban areas (red) versus missing from urban areas (teal) in each zone. Bird species were more specialized (i.e. narrower mean habitat breadth) toward lower latitudes (compare black mean lines among latitudinal zones). Species absent from urban areas were more specialized than those found in urban areas (compare teal vs. red points within each latitudinal zone; mean ± 95% CI), and the difference in specialization between urban and non-urban birds was greatest in the tropics. Polygons show smoothed distribution of raw species’ habitat breadths for each latitude x urbanization combination; statistical results in Table S3.

Our results also reveal an underappreciated effect of seasonality on both latitudinal and urbanization effects on diversity. As predicted, given widespread migration to high latitudes during summer breeding seasons, the avian latitudinal diversity gradient was less than half as steep in summer than winter, across urbanization categories (Fig. 3; Table S5). This result was robust to modelling decisions regarding the spatial scale of analyses, definition of ‘natural’, or minimum sampling thresholds (Tables S6 to S8). Indeed, the combined weakening effects of summer migration and urbanization appeared to erase the latitudinal diversity gradient, such that during summer urban bird richness did not increase discernibly toward the tropics (slope not different from 0, P = 0.97; Fig. 3). Seasonality also modified the negative effect of urbanization. In winter, urbanization no longer had a detectable negative effect on bird diversity at high latitudes (Fig. 3), consistent with hypotheses that birds are less sensitive to urbanization in high-latitude winter conditions (*32, 33*).

**Fig. 3.**
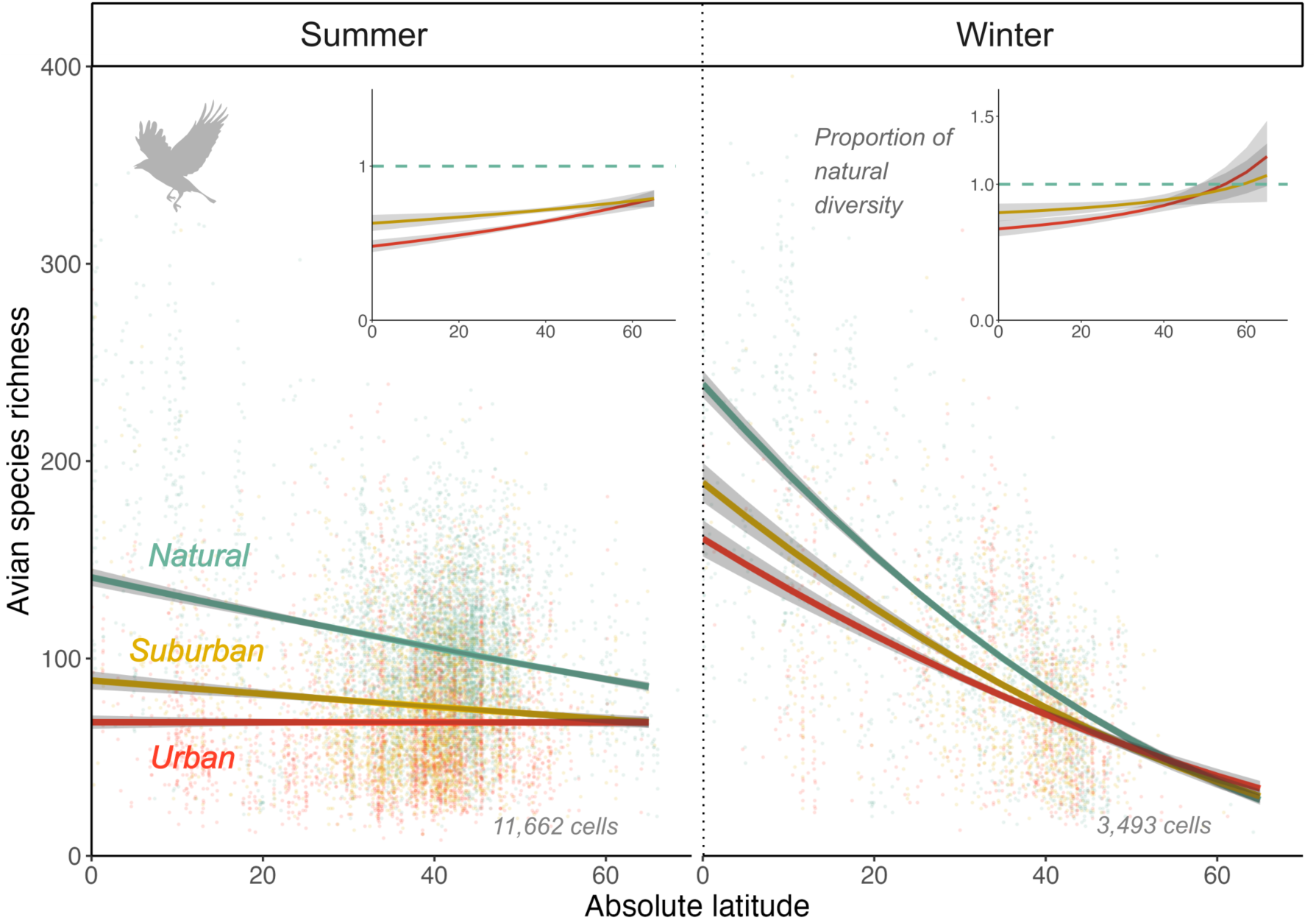
Effects of latitude and urbanization on diversity varied between seasons. Diversity in each 1×1 km cell was calculated independently for summer (June to August in the northern hemisphere, December to February in the southern hemisphere) and winter (June to August in the southern hemisphere, December to February in the northern hemisphere). Lines, grey polygons and points show marginal mean trend lines, 95% CI, and partial residuals extracted from a single model in which the season x latitude x urbanization interaction was significant (Table S5).

As macroecological patterns can differ between hemispheres (*34*), we explored whether the effect of latitude and urbanization differed between the northern and southern hemisphere. In our data, the latitudinal diversity gradient was steeper in the southern hemisphere in both natural and urban environments (Fig. S1; Table S1). Consistent with previous findings that fewer bird species migrate in the southern hemisphere (*30*), the latitudinal diversity gradient only differed seasonally in the northern hemisphere (Fig. S1). However, the lack of seasonal effect in the southern hemisphere should be interpreted with caution, as low winter sampling in the southern hemisphere (only 151 1×1 km cells had sufficient data for inclusion) may result in under- estimating winter diversity gradients.

The most parsimonious interpretation of our results is that urbanization has weakened the latitudinal diversity gradient in birds, driven at least partly by the disproportionate loss of more specialized species from urban areas. An alternate explanation for the first result could be that cities have been built in areas that happen to have a naturally weaker latitudinal diversity gradient, as cities are non-randomly distributed across landscapes (*35*) and it is plausible that the latitudinal diversity gradient varies among ecosystems. However, even if we only include natural cells within 20 km of an urban cell, natural areas still have a steeper diversity gradient than urban areas (Table S9; Fig. S7). Further, even among cells with <300 human inhabitants, more human-modified habitats have a weaker diversity gradient (Fig. S3). Thus the weaker gradient in urban areas does not seem to be an artefact of natural cells simply sampling different ecosystems than urban cells. Our results are also unlikely to be artefacts of geographic bias in sampling effort.

We find lower diversity in urban and suburban areas despite higher sampling effort in those cells (*36*), and lower diversity in the global north vs. tropics, despite greater sampling in wealthy northern countries (*37*). Thus our main results are conservative to sampling bias, and become more pronounced when stricter sampling thresholds are applied (Fig. S4). Finally, while we show that specialist species are disproportionately missing from cities (Fig. 3), we acknowledge that other factors may also contribute to the disproportionate loss of species from tropical cities. For example, cities tend to be larger and more intensively-urbanized toward lower latitudes (*38*) so may exclude a higher proportion of generalist species as well.

Our analyses of global avian assemblages show that urbanization has weakened the latitudinal diversity gradient, one of the most fundamental macroecological patterns on the planet. This puts urbanization on-par with other human impacts that have affected macroecological patterns at the global scale, such as large mammal declines (*39*), exotic species (*40*), and overfishing (*41*).

Further, our results provide global support for the long-standing hypotheses that tropical communities have more specialized species (*20*), the more recent hypothesis that specialists are particularly vulnerable to urbanization (*42*), and the new hypothesis that these features combine to weaken the latitudinal diversity gradient in cities (Fig. 1A). That disproportionate loss of specialists from tropical cities weakens the latitudinal diversity gradient also provides tantalizing support for the role of increased specialization in driving natural latitudinal patterns in diversity.

## Acknowledgments

We thank Ben Freeman for comments on an early version of the manuscript, Laura Pollock and Guillaume Larocque for advice on statistical analyses, and all of the birders who contribute their data to eBird and make this type of macroanalyses possible.

## Author contributions

All authors contributed to all stages and components of the paper.

## Data and materials availability

Raw eBird, urbanization, and specialization data used in analyses should be downloaded directly from their respective databases. Processed data and all code for analyses and visualization will be made publicly available upon acceptance in the Borealis data repository.

## Materials and Methods

### Bird occurrence data

We estimated bird species richness around the globe using the eBird basic dataset (version ebd_relMar-2023) (*43*). We used the R package ‘auk’ (*44*) to subset the data by the following criteria: 1) checklists were “complete”, meaning the observer recorded all bird species seen during their outing; 2) checklists were from outings where birding was the primary activity (i.e. followed the “stationary” or “traveling” protocols); 3) the distance traveled during the outing was less than 5 km so that the geolocated checklist point was not far from a given bird sighting; 4) the duration of the checklist was less than 5 h to reduce variability in sampling effort between checklists. We included checklists recorded between 2017 and 2022 inclusive to limit the bird sightings to a time period relevant for the urbanization data.

### Urbanization classes

We used a grid that divided the globe into 1×1 km cells, and classified each cell according to its degree of urbanization. To identify urban and suburban cells, we used the Settlement Model Layer (GHS-SMOD) of the 2020 Global Human Settlement data (GHS) developed by the European Commission (*45*). The GHS classifies urbanization based on the human population size and density in and around each grid cell (*46*). We considered ‘urban’ cells as those with at least 1500 inhabitants per km^2^ (GHS ‘urban centre’ category), and suburban cells as those with 300 to 1499 inhabitants (GHS ‘urban cluster’ category). To identify ‘natural’ cells, we selected cells with fewer than 300 inhabitants (GHS ‘rural’ category) that also had low habitat modification according to the Global Human Modification layer (GHM score < 0.5, from the NASA Socioeconomic Data and Applications center). The GHM gives each 1×1 km cell a cumulative human modification score from 0 (none) to 1 (highest) (*47*). To test how the choice of modification threshold affected results, we also ran analyses using modification thresholds of 0.25 and 0.75 to define natural cells (Fig. S3). We removed all cells classified as water or whose urbanization score was missing from either database.

### Species richness

We estimated species richness by aggregating eBird checklists into the 1×1 km raster cells of the urbanization layer and calculating the total number of species observed in each cell. As the world’s most poleward cities are at 70° N (Hammerfest, Norway) and 55° S (Ushuaia, Argentina), we removed natural cells beyond these latitudes.

To minimize the influence of sampling effort on species richness estimates, we set a minimum threshold for the number of checklists each cell needed to be included in analyses. To ensure our threshold was high enough to capture most of the diversity in even the richest cells, we determined how many checklists were needed to reach a sampling coverage of 95% in the 500 richest cells (‘iNEXT’ R package) (*48, 49*). Sample coverage is related to species accumulation curves, such that a 95% sampling coverage means there is only a 5% chance that an additional sample will yield a new species. We assigned each species a 1 or 0 based on whether it was recorded in a given checklist (presence = 1), summed frequencies over all checklists in the cell, and estimated coverage from the relative frequency of incidences. Doing this for each of the 500 richest cells yielded 500 estimates of the number of checklists needed to reach 95% coverage.

We took the 95^th^ percentile of this distribution, which gave a threshold of 102 checklists per cell. We therefore removed all cells with fewer than 102 checklists. This yielded 44,233 cells with both a measure of species richness and an urbanization score: 20,218 ‘natural’ cells, 13,987 ‘suburban’ cells, and 10,028 ‘urban’ cells (Fig. 1). We tested whether results were sensitive to the choice of minimum threshold by calculating the number of checklists needed to reach 90% and 98% sample coverage, taking the 95th percentile of these distributions, and re-running analyses of latitudinal diversity trends with these lower and higher thresholds (Fig. S4).

### Analyses of year-round diversity

To examine the effect of urbanization on the latitudinal diversity gradient, we fit a linear model with species richness per 1×1 km as the response and absolute latitude, urbanization category, and their interaction as predictors. Species richness was square-root transformed to meet assumptions of normality. We included a predictor for hemisphere (north or south) and the 3- way latitude x urbanization x hemisphere interaction to test whether latitudinal and urbanization effects on diversity differ between hemispheres. As diversity varies with elevation and precipitation (*50*), we included these as non-interacting covariates, using the elevation of each cell’s midpoint (‘elevatr’ package) (*51*) and mean annual precipitation from WorldClim (*52*).

Precipitation was removed from final models as it was never significant (F = 2.69, P = 0.10 in main analysis). Although we only included cells with more than 102 eBird checklists, we included the covariate log(number of checklists) to account for any remaining effect of sampling effort on estimated species richness. The model included a residual autocovariate as a fixed effect to account for spatial autocorrelation, with the neighborhood distance set to 100 km (chosen based on AIC scores) and weighted by inverse distance (*53*), a method of accounting for spatial autocorrelation used in other studies on latitudinal diversity gradients (*54*). Thus the model was: sqrt(species richness) ∼ absolute latitude x urbanization x hemisphere + elevation + log(number of checklists) + spatial covariate. To visualize model results, we used the ‘marginaleffects’ package (*55*) to calculate marginal effects of each predictor averaged over other predictor variables. The most poleward urban cells in our analyses were 65° latitude, so all reported predictions are from 0 to 65°.

### Specialization

To test how specialization varies with latitude and urbanization, we obtained species-level data on habitat and diet breadth. For habitat breadth, we used an existing index that combines the number of IUCN habitat categories in which each species occurs and the diversity of other taxa with which it co-occurs (i.e. a generalist occurs in habitats that vary considerably in species composition (*56*)). We derived an estimate of diet breadth using the EltonTraits database (*57*), which contains the proportion of a species’ diet that come from ten diet classes. From these we calculated the Gini index, which measures inequality and has previously been used as a measure of specialization in birds (*58, 59*). The Gini index ranges from 0 (equal percentage of diet in each class, i.e. maximum generalization) to 1 (100% of diet in one class, i.e. maximum specialization). We then calculated diet breadth for each species as 1 – the Gini index, so that larger values represented greater diet breadth. Of 10,163 bird species in our data, we were able to estimate habitat breadth for 7,941 and diet breadth for 6,580 species.

We tested predictions that lower latitudes have more specialized species (*20*), that urbanization disproportionately excludes more specialized species (*42*), and that these factors combined cause disproportionate species losses from lower vs. higher latitude cities (Fig. 1A). To compare the relative number of specialists at different latitudes, we binned cells into four latitudinal zones based on the absolute latitude of the center of the cell: tropical (0° to 23.44°), subtropical (>23.44° to 35°), temperate (>35° to 50°) and subpolar (>50° to 70°; cells above 70° were excluded due to lack of urban areas, see ‘*Species richness*’). For each bird species in each latitudinal zone, we assigned an urban presence score: ‘urban present’ if the species was ever found in urban cells in that zone, ‘urban absent’ if the species was never found in urban cells in that zone. A species could be ‘urban absent’ in one latitudinal zone but ‘urban present’ in another. We ran one linear model per specialization response (habitat or diet breadth, log transformed to meet assumptions of normality), with urban presence, latitudinal zone, and their interaction as predictors. We used the ‘marginaleffects’ package to determine whether mean specialization differed between latitudes and between urban-absent and urban-present birds.

### Seasonal diversity

To explore whether the effects of latitude, urbanization, and specialization differ between seasons, we filtered the original eBird data for checklists recorded during summer (June to August in the northern hemisphere, December to February in the southern hemisphere) or winter (December to February in the northern hemisphere, June to August in the southern hemisphere). We then calculated summer and winter species richness independently for each cell. We reduced the threshold number of checklists for inclusion, as few cells had 102 checklists in a 3-month window, and because fewer checklists are needed to reach 95% sampling coverage of 3-month diversity than year-round diversity in the 500 richest cells). Instead of using the 95th percentile of the distribution needed to reach 95% sampling coverage of year-round diversity (102 checklists; main analyses), we used the 75th percentile. This gave a threshold of 68 checklists, so we removed cells with fewer than 68 checklists in a given season. While this lower threshold may underestimate diversity in the richest cells (low latitudes and natural areas, particularly low latitudes during winter, Fig. 3), the patterns we note would be even stronger with increased sampling coverage (e.g. steeper latitudinal diversity gradients in natural areas, especially in winter). This yielded 11,662 cells for summer richness (5,436 natural, 3,447 suburban, 2,779 urban) and 3,493 cells for winter richness (1589 natural, 1069 suburban, 835 urban).

We re-ran both the diversity and specialization analyses including only the summer and winter data, and tested whether results differed between seasons. For diversity, we ran the same linear model described for year-round data (Table S1), but with a 4-way interaction between latitude x urbanization x hemisphere x season to test whether 1) the latitudinal diversity gradient differed between seasons (latitude x season), and 2) the effect of urbanization on the latitudinal diversity gradient differed between seasons (latitude x urbanization x season). Again, we accounted for spatial autocorrelation by including a residual autocovariate term to account for spatial autocorrelation, with the neighborhood distance set to 500 km (chosen based on AIC scores) and weighted by inverse distance. For the specialization analysis, we re-ran the linear model with the 3-way interaction urban presence x latitudinal zone x season to test whether the effect of urbanization on specialists differed among seasons.

### Sensitivity to scale

Because species-area relationships are non-linear, we tested whether the diversity trends we detected were sensitive to spatial scale by redoing analyses with both urbanization and bird diversity data grouped into 5×5 km cells, rather than the 1×1 km cells in our main analyses. We could not use a smaller scale as urbanization data are only available at 1×1 resolutions, and we did not test a larger scale as too few cells could have been classified as natural or urban for robust analyses of global trends. We assigned each 5×5 km cell an urbanization classification if it contained 70% or more of a given category (i.e. at least 18 of the 1×1 km cells shared the same urbanization category). For year-round data, we recalculated the sampling intensity needed to reach 95% coverage in the 500 richest 5×5 km cells and used the 95th percentile of that distribution, which yielded a sampling intensity threshold of 163 checklists. Because there are fewer 5×5 vs 1×1 km cells and because all cells with <70% in any urbanization category were removed, sample sizes were significantly reduced; 1172 natural cells, 44 suburban cells, and 421 urban cells. As a result, full models did not converge properly, so we removed the interaction terms involving hemisphere, which were not significant in our main analyses. The same logical process for analyses of seasonal data yielded a sampling intensity threshold of 84 checklists. We re-ran models of year-round diversity and seasonal diversity; results are summarized in Table S6 visualized in Fig. S2.

## Supplementary Figures

**Fig. S1.**
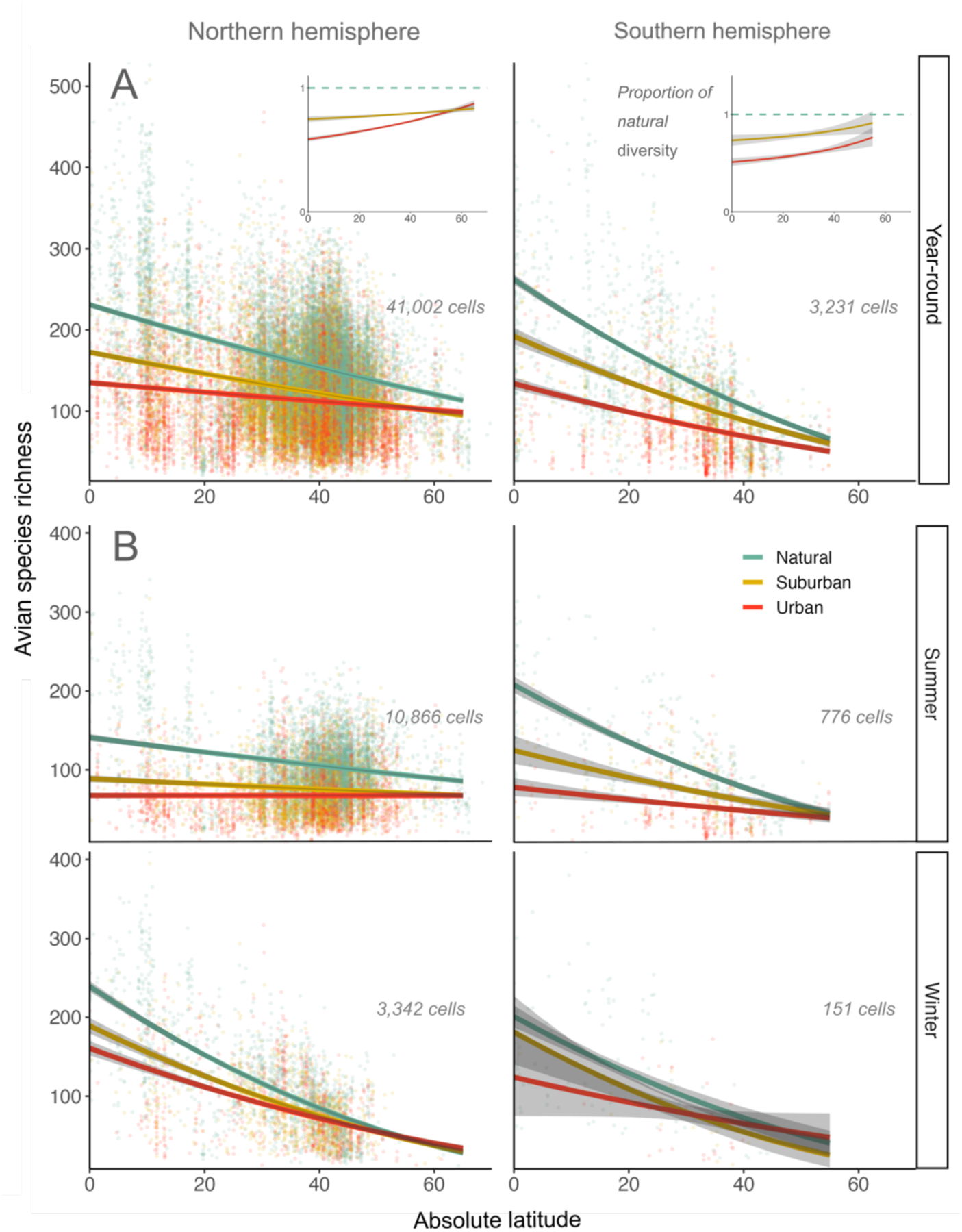
Hemisphere-specific effects of urbanization on the avian latitudinal diversity gradient (LDG). **A**) The significant latitude x urbanization x hemisphere interaction from our main model (Table S1). While the slopes of the LDG differ among hemispheres, in both hemispheres urbanization lowers diversity and weakens the LDG by disproportionately reducing diversity at low latitudes, consistent with the ‘*Disproportionate loss from tropics*’ hypothesis (H3; Fig. 1A). **B**) The non-significant latitude x urbanization x hemisphere x season interaction from the seasonal model (F=12.37, P = 0.09; Table S5). Urbanization weakens the LDG in all season x hemisphere combinations, but the effect of season on the LDG (stronger in winter than summer; Fig. 3) is only apparent in the northern hemisphere (though note the small amount of seasonal data available for the southern hemisphere). Lines, grey polygons, and points show model marginal mean trend lines, 95% confidence intervals, and partial residuals, respectively. Confidence intervals are sometimes narrower than the plotted trend line so are hard to see. Predictions extend to 65° N but only 55° S as there are no urbanized areas south of 55° S.

**Fig. S2.**
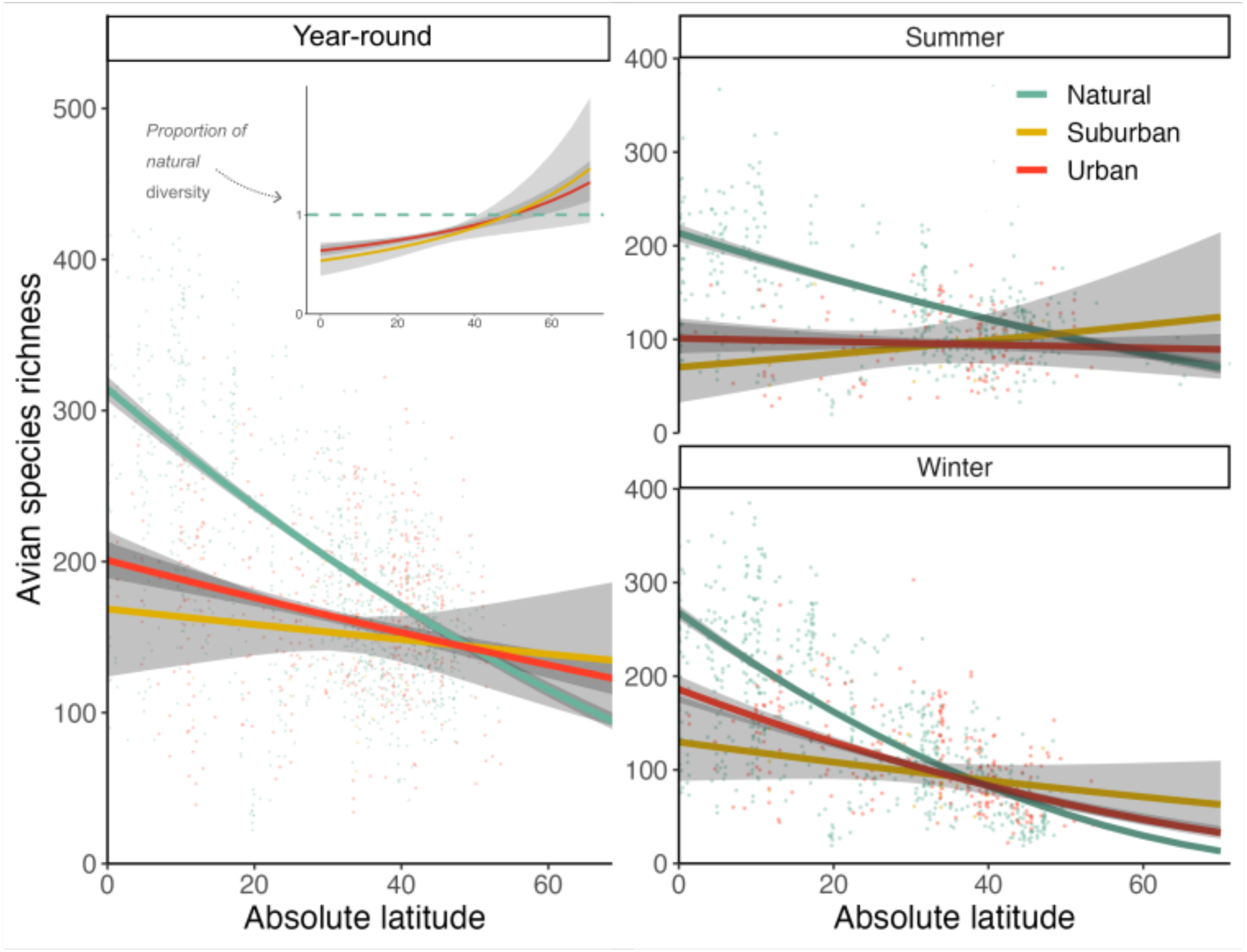
Main analyses redone with data compiled at the 5×5 km scale. Results are broadly consistent with results from main models using 1×1 km grid cells (Fig. 1C & Fig. 3), in that urbanization weakens the latitudinal diversity both year round (left) and in each season (right), and that the year-round diversity gradient is stronger in winter than summer. However, using a 5×5 km scale reduced the number of cells by more than 5-fold, so the results for suburban cells are no longer significantly different than those of urban cells. Lines, shaded polygons, and points show model trend lines, 95% confidence intervals, and partial residuals, respectively, taken from models summarized in Table S6. Values are marginal means, i.e. averaged across values of other model predictors, including hemisphere.

**Figure S3.**
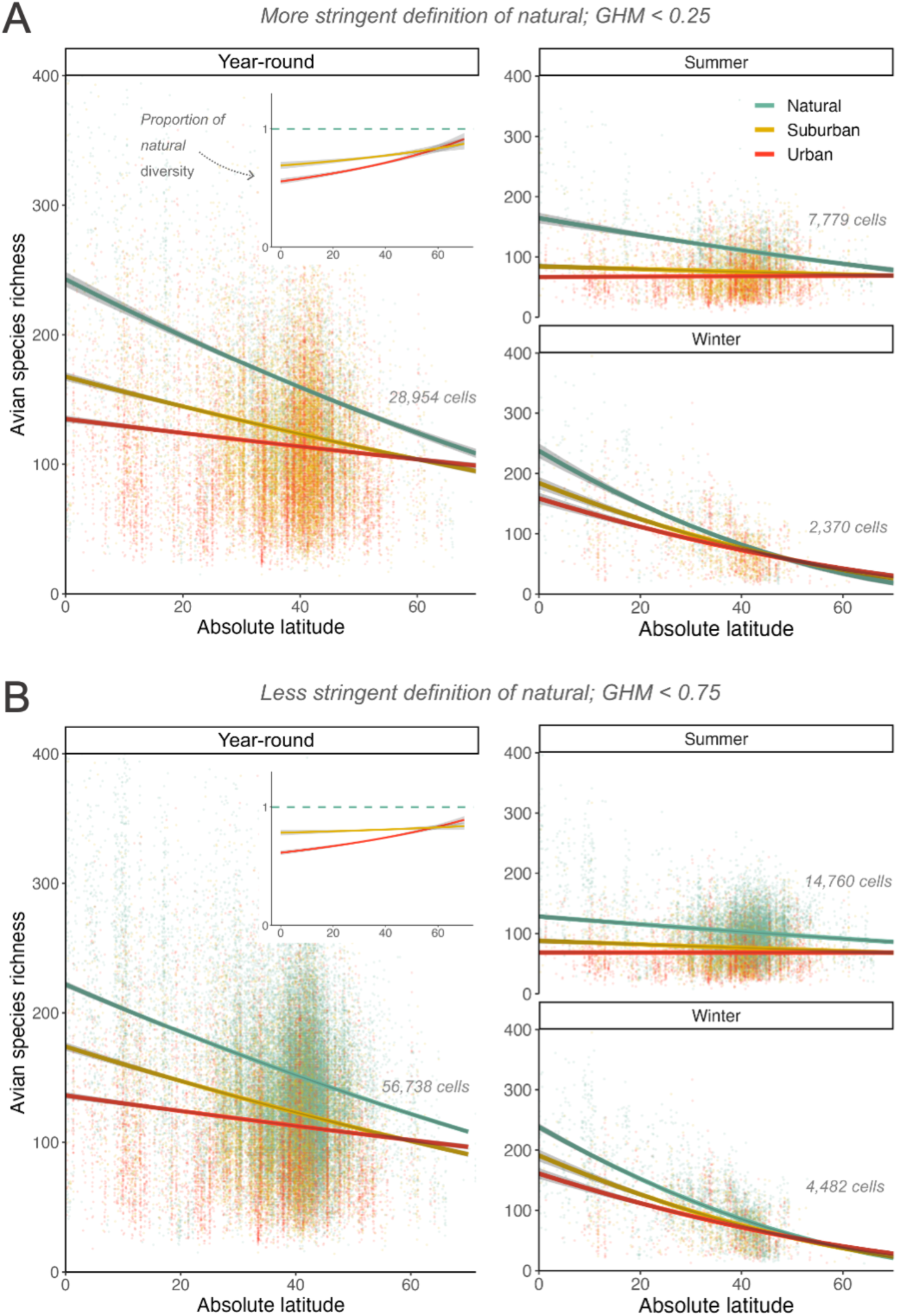
Higher thresholds for ‘natural’ steepen the latitudinal diversity gradient but do not alter main conclusions. Main results classified any cell with fewer than 300 human inhabitants (i.e. ‘rural’) as ‘natural’ if it had a Global Human Modification score of 0.5 or less (Fig. 1C & Fig. 3). **A**) Requiring even less human modification for natural cells (GHM 0.25 or less) decreased our sample size (n = 4,937) and yielded a steeper latitudinal diversity gradient in natural cells than in our original analyses (124% increase from 70° to 0°, compared to the 118% increase in Fig. 1C). **B**) Using a more permissive threshold (GHM of 0.75 or less) increased sample size (n = 32,721 natural cells) and yielded a weaker diversity gradient. Thus increased habitat modification even in areas with low human populations weakened the latitudinal diversity gradient in birds. Figure formatting as in Fig. S1; statistical results in Table S7.

**Fig. S4.**
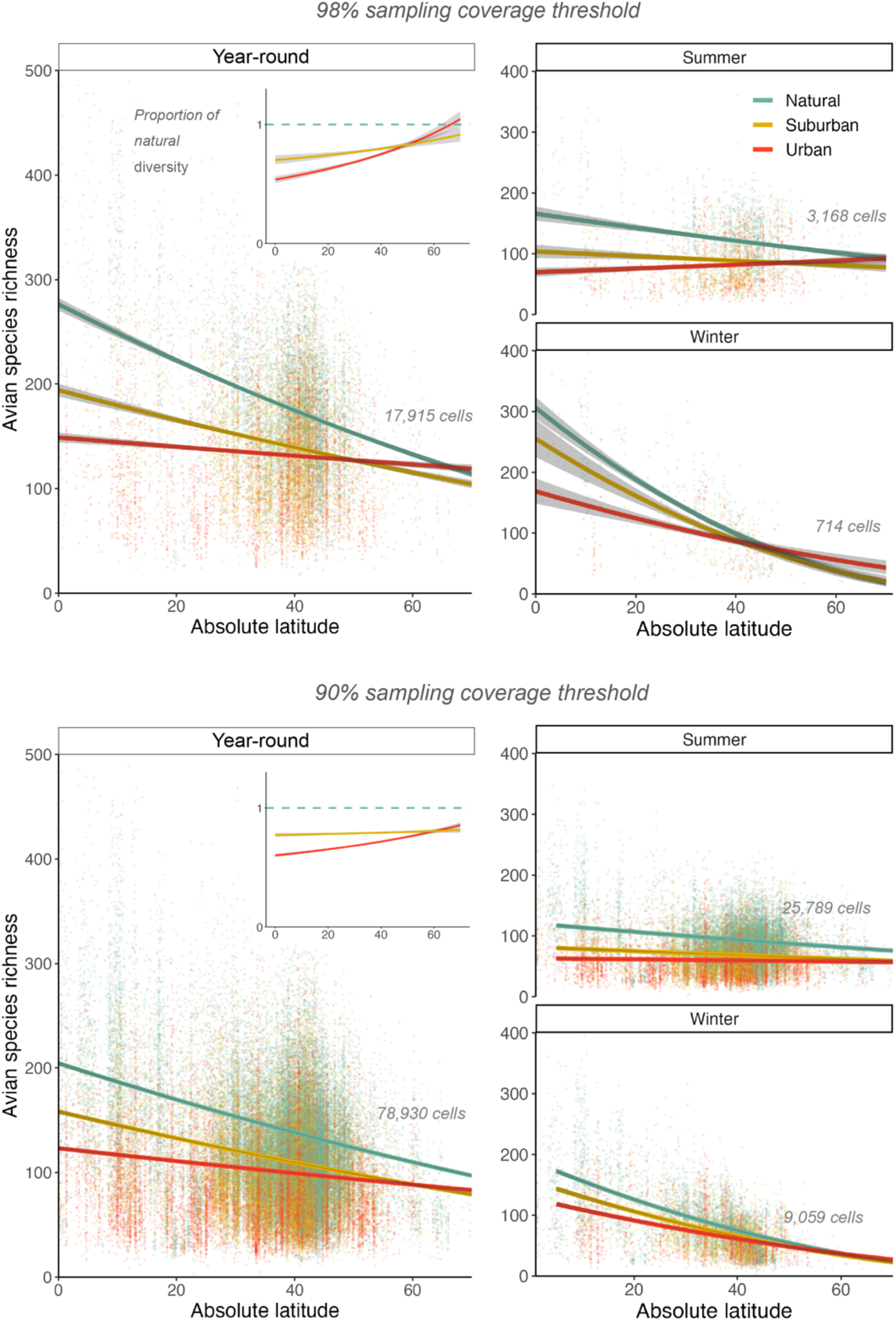
Higher thresholds for sampling coverage increased detected diversity but did not change main conclusions. Sampling intensity threshold in main analyses was determined using the estimated number of eBird checklists needed to reach a 95% sampling coverage in the 500 richest cells; analyses of year-round diversity used the 95th percentile of this distribution (102 checklists per cell), while analyses of seasonal diversity used the 75th percentile (68 checklists per cell in a given season). Main conclusions remain unchanged if we instead use the 95th and 75th percentile of the distribution needed to reach 98% sampling coverage (i.e. a stricter threshold; 258 and 174 checklists; top panels), or 90% sampling coverage (51 and 33 checklists; bottom panels). Requiring greater sampling coverage reduces the number of cells in analyses but increases the mean detected bird diversity, particularly in high-diversity areas (low-latitude and natural areas). This results in a steeper detected latitudinal diversity gradient in natural areas and a greater difference in diversity gradient between natural and urban areas, providing even stronger support for our main conclusion. With 95% sampling coverage, mean year-round bird diversity at 70° and 0° is 106 and 230 species/km^2^ in natural areas (118% increase from 70° to equator), and 96 and 118 species/km^2^ in urban areas (22% increase; Fig. 1C). With 98% sampling coverage, mean diversity at 70° and 0° increases to 119 and 276 species/km^2^ in natural areas (131% increase), and 119 and 149 species/km^2^ in urban areas (25% increase). Lowering the threshold has the opposite effects. For all three sampling thresholds, the latitudinal diversity gradient in birds is weaker in urban than suburban than natural areas, and is weaker in summer than winter. Figure formatting as in Fig. S1; statistical results in Table S8.

**Fig. S5.**
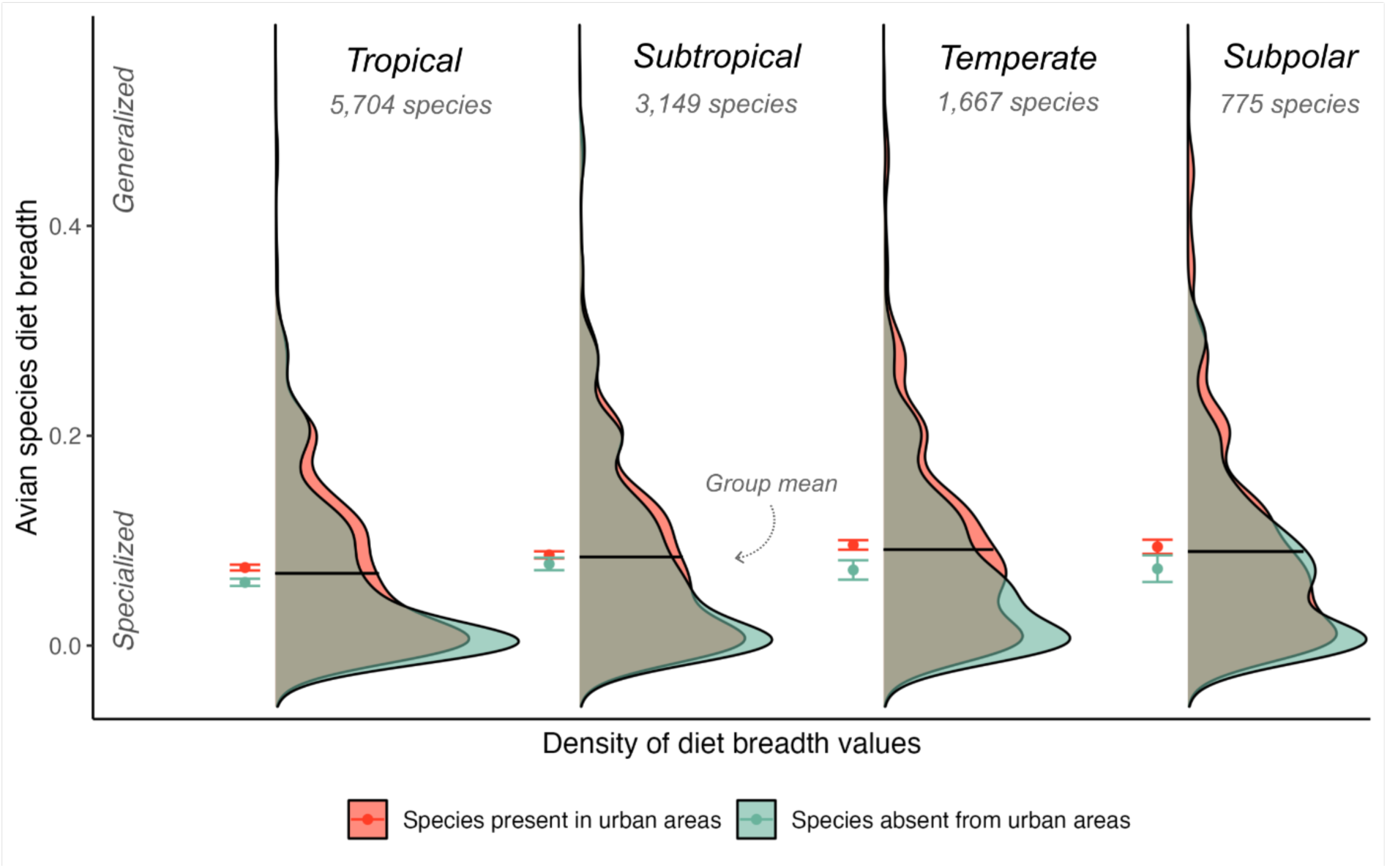
Diet specialization vs. latitude and urbanization. Distribution of diet-breadth scores in each of four latitudinal zones for bird species present in urban areas (red) and bird species absent from urban areas (teal) in each zone. Bird species were more specialized (i.e. narrower mean diet breadth) toward lower latitudes (compare black median lines among latitudinal zones), and this trend was driven by species that could be found in urban areas (red points; mean ± SE). Species absent from urban areas tended to be more specialized than those found in urban areas in each latitudinal zone (compare teal vs. red points within each latitudinal zone), though the difference was not always significant. Statistical results in Table S3.

**Fig. S6.**
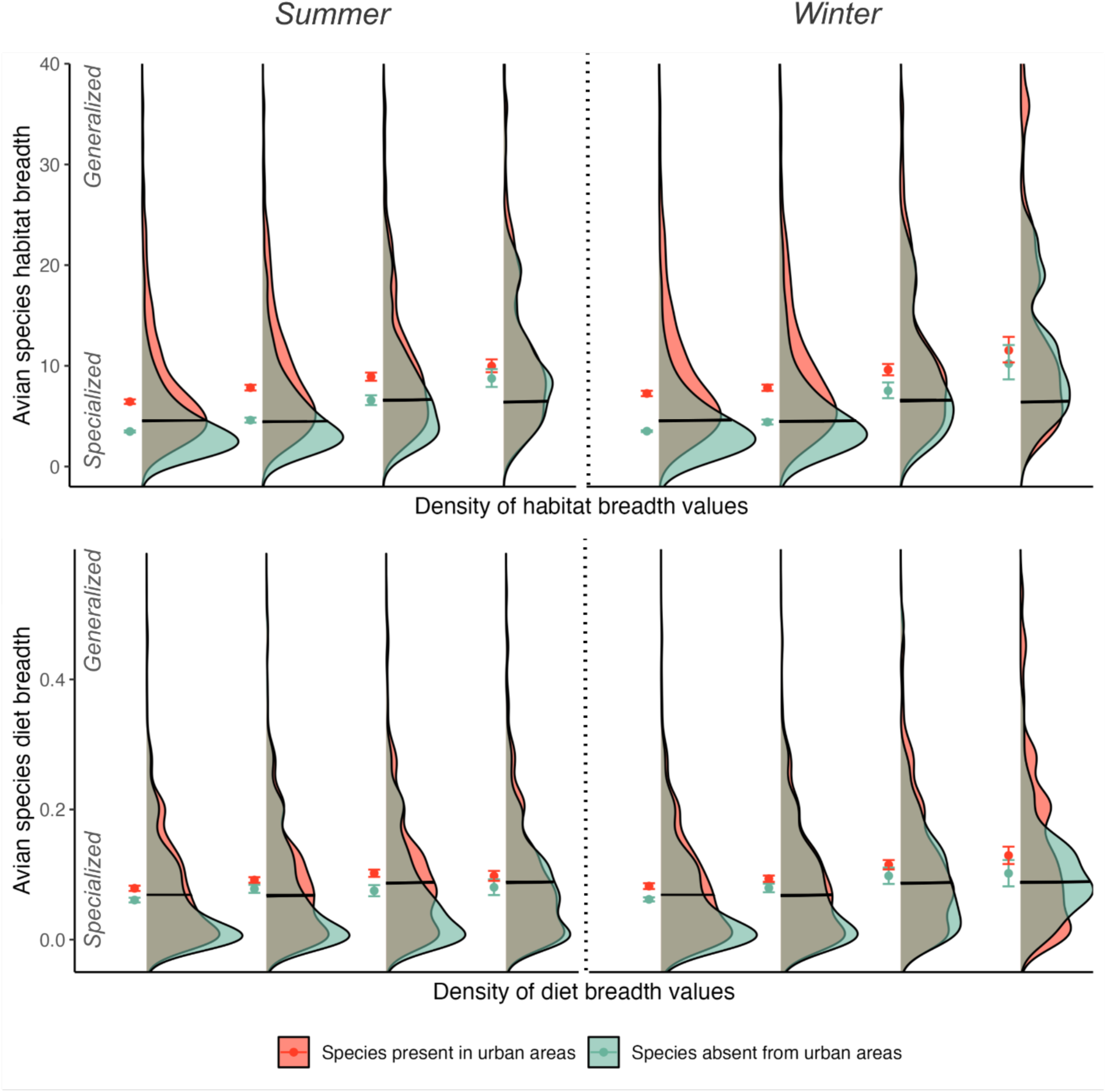
Season-specific effects of urbanization and latitude on specialization. Formatting as in Figures 2 and S5, statistical results in Table S4. Birds had more specialized habitats and diets from subpolar to tropical latitudes in both seasons (compare black median lines). Birds that could be found in urban areas were always less specialized on average than those that were absent from urban areas in that latitudinal zone, though the difference was not always significant (compare red vs teal points; mean ± 95% CI).

**Fig. S7.**
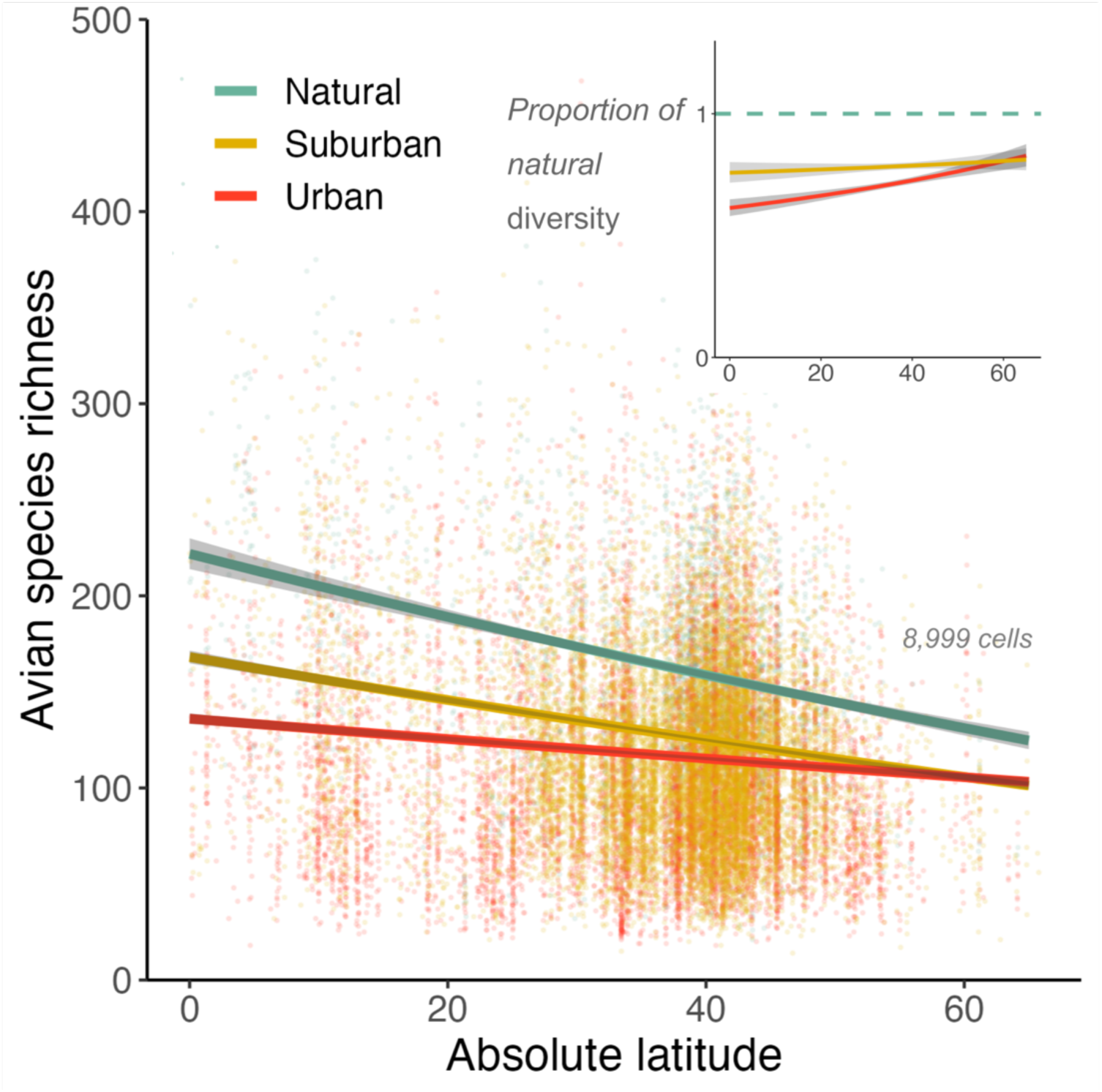
Even natural areas close to cities have a steeper latitudinal diversity gradient than urban areas. To examine whether the weaker latitudinal diversity gradient detected in urban areas might be an artefact of urban and natural cells sampling different ecosystems, we re-ran our main diversity analysis including only natural cells that were within 20 km of an urban cell, such that natural and urban cells presumably sample similar regional bird communities. This reduced our sample size of natural cells from 20,218 cells (Fig. 1) to 2902 cells (above). Despite this much lower sample size, and the fact that natural cells may well experience ecological effects of cities so nearby, natural areas still have significantly higher bird diversity and a steeper avian latitudinal diversity gradient than suburban or urban areas. Figure formatting as in Fig S1, statistical results in Table S9.

## Supplementary Tables

**Table S1.** Statistical results for main analysis of how bird diversity varies with latitude and urbanization. Response is species richness per 1×1 km grid cell, including only cells with at least 120 complete eBird checklists. Predictors are: absolute latitude; urbanization (natural, suburban, urban); hemisphere (north or south); number of eBird checklists per cell to account for sampling effort; and elevation of each cell’s midpoint. The model included an inverse weighted spatial autocorrelation term with a 100 km neighborhood distance chosen based on AIC scores. The effects of latitude and urbanization are plotted in Fig. 1C (across values of other predictors).

| **Linear model: how urbanization affects year-round bird diversity across latitudes**<br>$\sqrt{\text{species richness}} \sim \text{absolute latitude} \times \text{urbanization} \times \text{hemisphere} +$<br>$\log(\text{number of checklists}) + \text{elevation} + \text{spatial autocovariate}$<br>(df = 14 and 44,218, $R^2 = 0.53$) | | | | |
| --- | --- | --- | --- | --- |
| **Type III ANOVA Results** |  |  |  |  |
| **Variable** | **Sum Sq** | **Df** | **F value** | **P value** |
| Absolute latitude | 9184.42 | 1 | 3230.14 | < 2E-16 |
| Urbanization | 6839.25 | 2 | 1202.67 | < 2E-16 |
| Hemisphere | 505.91 | 1 | 177.93 | < 2E-16 |
| Log(number of checklists) | 42764.15 | 1 | 15040.05 | < 2E-16 |
| Elevation | 1315.34 | 1 | 462.60 | < 2E-16 |
| Spatial autocovariate | 49703.62 | 1 | 17480.64 | < 2E-16 |
| Absolute latitude x urbanization | 1512.55 | 2 | 265.98 | < 2E-16 |
| Absolute latitude x hemisphere | 1420.81 | 1 | 499.69 | < 2E-16 |
| Urbanization x hemisphere | 88.26 | 2 | 15.52 | 0.000000183 |
| Absolute latitude x urbanization x hemisphere | 43.82 | 2 | 7.71 | 0.000451 |
| **Regression coefficients** |  |  |  |  |
| **Variable** | **Estimate** | **Std. Error** | **T value** | **P value** |
| Intercept | 7.64 | 0.077 | 99.63 | < 2E-16 |
| Absolute latitude | -0.07 | 0.0012 | -56.83 | < 2E-16 |
| Urbanization (Suburban) | -2.06 | 0.079 | -26.20 | < 2E-16 |
| Urbanization (Urban) | -3.57 | 0.073 | -26.20 | < 2E-16 |
| Hemisphere (Southern) | 1.49 | 0.11 | 13.33 | < 2E-16 |
| Log(number of checklists) | 1.29 | 0.011 | 122.64 | < 2E-16 |
| Elevation | -0.00035 | 0.000016 | -21.51 | < 2E-16 |
| Spatial autocovariate | 1.007 | 0.002 | 8.74 | < 2E-16 |
| Absolute latitude x urbanization (suburban) | 0.018 | 0.002 | 8.74 | < 2E-16 |
| Absolute latitude x urbanization (urban) | 0.044 | 0.002 | 21.06 | < 2E-16 |
| Absolute latitude x hemisphere (southern) | -0.08 | 0.0036 | -22.35 | < 2E-16 |
| Urbanization (suburban) x hemisphere (southern) | -0.33 | 0.23 | -1.44 | 0.15 |
| Urbanization (urban) x hemisphere (southern) | -1.18 | 0.21 | -5.57 | 0.0000000254 |
| Absolute latitude x urbanization (suburban) x hemisphere (southern) | 0.019 | 0.007 | 2.67 | 0.0076 |
| Absolute latitude x urbanization (urban) x hemisphere (southern) | 0.023 | 0.0066 | 3.51 | 0.00046 |

**Table S2.** Predicted marginal-mean species richness shown in Fig. 1 at every 10° of absolute latitude, derived from the linear model in Table S1. Year-round bird richness is estimated for the median value of elevation and log(number of checklists) and across hemispheres (though weighted more heavily toward the northern hemisphere given the greater amount of data there). From 75 to 0° latitude, natural cells gain an average of 118 species whereas urban areas gain only 36 species, yielding an overall diversity gradient more than 3-times stronger in natural vs. urban areas.

| Absolute latitude | Urbanization category | Predicted bird richness (mean # of species) | Low CI | High CI |
| --- | --- | --- | --- | --- |
| 0 | Natural | 230.94 | 228.04 | 233.85 |
| 0 | Suburban | 172.57 | 169.4 | 175.78 |
| 0 | Urban | 135.13 | 132.61 | 137.69 |
| 5 | Natural | 220.41 | 217.92 | 222.92 |
| 5 | Suburban | 165.76 | 163.03 | 168.5 |
| 5 | Urban | 132.15 | 129.96 | 134.36 |
| 10 | Natural | 210.13 | 208.03 | 212.25 |
| 10 | Suburban | 159.08 | 156.78 | 161.39 |
| 10 | Urban | 129.19 | 127.33 | 131.07 |
| 15 | Natural | 200.1 | 198.36 | 201.84 |
| 15 | Suburban | 152.54 | 150.65 | 154.43 |
| 15 | Urban | 126.27 | 124.71 | 127.84 |
| 20 | Natural | 190.31 | 188.91 | 191.71 |
| 20 | Suburban | 146.13 | 144.62 | 147.64 |
| 20 | Urban | 123.38 | 122.11 | 124.67 |
| 25 | Natural | 180.76 | 179.66 | 181.87 |
| 25 | Suburban | 139.86 | 138.7 | 141.03 |
| 25 | Urban | 120.53 | 119.5 | 121.56 |
| 30 | Natural | 171.47 | 170.61 | 172.32 |
| 30 | Suburban | 133.73 | 132.86 | 134.61 |
| 30 | Urban | 117.71 | 116.86 | 118.56 |
| 35 | Natural | 162.41 | 161.72 | 163.1 |
| 35 | Suburban | 127.74 | 127.05 | 128.43 |
| 35 | Urban | 114.92 | 114.16 | 115.68 |
| 40 | Natural | 153.61 | 152.95 | 154.26 |
| 40 | Suburban | 121.88 | 121.21 | 122.56 |
| 40 | Urban | 112.17 | 111.37 | 112.96 |
| 45 | Natural | 145.04 | 144.3 | 145.79 |
| 45 | Suburban | 116.16 | 115.35 | 116.98 |
| 45 | Urban | 109.44 | 108.51 | 110.38 |
| 50 | Natural | 136.73 | 135.82 | 137.63 |
| 50 | Suburban | 110.58 | 109.55 | 111.62 |
| 50 | Urban | 106.76 | 105.63 | 107.89 |
| 55 | Natural | 128.66 | 127.56 | 129.76 |
| 55 | Suburban | 105.14 | 103.86 | 106.43 |
| 55 | Urban | 104.10 | 102.75 | 105.46 |
| 60 | Natural | 120.83 | 119.54 | 122.13 |
| 60 | Suburban | 99.83 | 98.31 | 101.37 |
| 60 | Urban | 101.48 | 99.89 | 103.08 |
| 65 | Natural | 113.25 | 111.77 | 114.75 |
| 65 | Suburban | 94.67 | 92.9 | 96.45 |
| 65 | Urban | 98.89 | 97.07 | 100.74 |

**Table S3.** Statistical results for how specialization varies with latitude and between species present in vs. absent from cities. To assess specialization, each bird species was assigned a continuous score for habitat breadth and diet breadth (lower breath score = more specialized). Within each latitudinal zone (tropical, subtropical, temperature, subpolar), every species was also classified as either ‘urban present’ if it was present in at least 1 urban cell in that zone (at any time of the year), otherwise ‘urban absent’. Models test whether mean specialization score varies among latitudinal zones and between species that were present in urban areas vs. absent from urban areas (Fig. 2).

| ANOVA: habitat breadth vs. latitude zone and urban presence<br>log(habitat breadth) ~ latitude zone x urban presence |  |  |  |  |  |
| --- | --- | --- | --- | --- | --- |
| Variable | Df | Sum sq. | Mean sq. | F value | P value |
| Latitude zone | 3 | 955 | 318.4 | 535.4 | < 0.0000001 |
| Urban presence | 1 | 787 | 786.7 | 1323.0 | < 0.0000001 |
| Latitude zone x urban presence | 3 | 43 | 14.5 | 24.4 | < 0.0000001 |
| ANOVA: diet breadth vs. latitude zone and urban presence<br>log(diet breadth) ~ latitude zone x urban presence |  |  |  |  |  |
| Variable | Df | Sum sq. | Mean sq. | F value | P value |
| Latitude zone | 3 | 0.87 | 0.29 | 48.0 | < 0.0000001 |
| Urban presence | 1 | 0.41 | 0.41 | 67.2 | < 0.0000001 |
| Latitude zone x urban presence | 3 | 0.04 | 0.013 | 2.2 | 0.08 |

**Table S4.** Season-specific effects of latitude and urbanization on specialization. Model responses are a species-specific value for habitat or diet breadth (lower breath score = more specialized). Data include only cells with at least 68 eBird checklists per season. Within each season and each latitudinal zone (tropical, subtropical, temperature, subpolar), every species was classified as either ‘urban present’ or ‘urban absent’. Models test whether season affects how mean specialization score varies among latitudinal zones and among bird species that were vs. were never found in urban areas during a given season. Model results are visualized in Fig. S6.

| ANOVA: habitat breadth vs. latitude zone and urban presence by season<br>log(habitat breadth) ~ latitude zone x urban presence x season |  |  |  |  |  |
| --- | --- | --- | --- | --- | --- |
| Variable | Df | Sum sq. | Mean sq. | F value | P value |
| Latitude zone | 3 | 1306 | 435.5 | 730.91 | < 0.0000001 |
| Urban presence | 1 | 1249 | 1249.4 | 2096.97 | < 0.0000001 |
| Season | 1 | 5 | 5.4 | 9.091 | 0.0026 |
| Latitude zone x urban presence | 3 | 103 | 34.3 | 57.63 | < 0.0000001 |
| Latitude zone x season | 3 | 8 | 2.6 | 4.34 | 0.0046 |
| Urban presence x season | 1 | 4 | 4.2 | 7.09 | 0.0078 |
| Latitude zone x urban presence x season | 3 | 3 | 1.1 | 1.77 | 0.15 |
| ANOVA: diet breadth vs. latitude zone and urban presence by season<br>log(diet breadth) ~ latitude zone x urbanization category x season |  |  |  |  |  |
| Variable | Df | Sum sq. | Mean sq. | F value | P value |
| Latitude zone | 3 | 2.14 | 0.71 | 116.07 | < 0.0000001 |
| Urban presence | 1 | 0.97 | 9.97 | 157.45 | < 0.0000001 |
| Season | 1 | 0.08 | 0.075 | 12.25 | 0.00047 |
| Latitude zone x urban presence | 3 | 0.03 | 0.094 | 1.52 | 0.21 |
| Latitude zone x season | 3 | 0.15 | 0.049 | 7.98 | 0.000026 |
| Urban presence x season | 1 | 0.00 | 0.0004 | 0.067 | 0.80 |
| Latitude zone x urban presence x season | 3 | 0.01 | 0.0041 | 0.67 | 0.57 |

**Table S5.** Season-specific effects of latitude and urbanization on bird diversity. Response is bird species richness per 1×1 km grid cell calculated for either summer months (July to Aug or Dec to Feb in northern and southern hemisphere, respectively), or winter months (July to Aug or Dec to Feb in southern and northern hemisphere, respectively). Data include only cells with at least 68 eBird checklists in that season. Predictors are those described in Table S1, plus a categorical predictor for season (summer or winter) and its interactions. The model included an inverse weighted spatial autocorrelation term with a 500 km neighborhood distance chosen based on AIC scores.

| **Linear model: how urbanization affects seasonal bird diversity across latitudes**<br>$\sqrt{\text{species richness}} \sim \text{absolute latitude} \times \text{urbanization} \times \text{hemisphere} \times \text{season} + \log(\text{number of checklists}) + \text{elevation} + \text{spatial autocovariate}$<br>(df = 26 and 15128 , $R^2 = 0.52$) | | | | |
| --- | --- | --- | --- | --- |
| Type III ANOVA Results |  |  |  |  |
| Variable | Sum Sq. | Df | F value | P value |
| Absolute latitude | 783.4 | 1 | 305.5 | < 2E-16 |
| Urbanization | 1788.7 | 2 | 348.8 | < 2E-16 |
| Hemisphere | 586.1 | 1 | 228.6 | < 2E-16 |
| Season | 1544.9 | 1 | 602.5 | < 2E-16 |
| Log(number of checklists) | 5773.8 | 1 | 2251.8 | < 2E-16 |
| Elevation | 930.7 | 1 | 363.0 | < 2E-16 |
| Spatial autocovariate | 13486.1 | 1 | 5259.6 | < 2E-16 |
| Absolute latitude x urbanization | 332.8 | 2 | 64.9 | < 2E-16 |
| Absolute latitude x hemisphere | 620.8 | 1 | 242.1 | < 2E-16 |
| Urbanization x hemisphere | 64.4 | 2 | 12.6 | < 2E-16 |
| Absolute latitude x season | 2139.3 | 1 | 834.3 | < 2E-16 |
| Urbanization x season | 37.9 | 2 | 7.4 | 0.00061 |
| Hemisphere x season | 311.4 | 1 | 121.4 | < 2E-16 |
| Absolute latitude x urbanization x hemisphere | 56.7 | 2 | 11.1 | 1.58E-05 |
| Absolute latitude x urbanization x season | 6.5 | 2 | 1.3 | 0.28 |
| Absolute latitude x hemisphere x season | 141.3 | 1 | 55.1 | 1.20E-13 |
| Urbanization x hemisphere x season | 11.4 | 2 | 2.2 | 0.11 |
| Absolute latitude x urbanization x hemisphere x season | 12.4 | 2 | 2.4 | 0.090 |

| Regression coefficients |  |  |  |  |
| --- | --- | --- | --- | --- |
| Variable | Estimate | SE | T value | P value |
| Intercept | 6.39 | 0.15 | 42.4 | < 2E-16 |
| Absolute latitude | -0.04 | 0.0023 | -17.5 | < 2E-16 |
| Urbanization (suburban) | -2.45 | 0.16 | -15.7 | < 2E-16 |
| Urbanization (urban) | -3.65 | 0.14 | -25.8 | < 2E-16 |
| Hemisphere (south) | 3.18 | 0.21 | 15.1 | < 2E-16 |
| Season (winter) | 3.58 | 0.15 | 24.6 | < 2E-16 |
| Log(number of checklists) | 1.11 | 0.023 | 47.4 | < 2E-16 |
| Spatial autocovariate | 1.04 | 0.014 | 72.5 | < 2E-16 |
| Elevation | 0.02 | 0.0039 | 5.6 | 1.9E-8 |
| Absolute latitude x urbanization (suburban) | 0.002 | 0.014 | 72.5 | < 2E-16 |
| Absolute latitude x urbanization (urban) | 0.04 | 0.014 | 72.5 | < 2E-16 |
| Absolute latitude x hemisphere (south) | -0.11 | 0.0066 | -15.6 | < 2E-16 |
| Urbanization (suburban) x hemisphere (south) | -0.86 | 0.48 | -1.8 | 0.075 |
| Urbanization (urban) x hemisphere (south) | -2.03 | 0.04 | -5.0 | 6.91E-7 |
| Absolute latitude x season (winter) | -0.12 | 0.004 | -28.9 | < 2E-16 |
| Urbanization (suburban) x season (winter) | 0.74 | 0.27 | 2.8 | 0.0051 |
| Urbanization (urban) x season (winter) | 0.87 | 0.26 | 3.4 | 0.0007 |
| Season (winter) x hemisphere (south) | -3.82 | 0.35 | -11.0 | < 2e-16 |
| Absolute latitude x urbanization (suburban) x hemisphere (south) | 0.04 | -0.15 | 2.4 | 0.014 |
| Absolute latitude x urbanization (urban) x hemisphere (south) | 0.06 | 0.012 | 4.5 | 7.44E-6 |
| Absolute latitude x urbanization (suburban) x season (winter) | 0.007 | 0.0072 | 1.5 | 0.13 |
| Absolute latitude x urbanization (urban) x season (winter) | 0.14 | 0.015 | 7.4 | 1.2E-13 |
| Absolute latitude x season (winter) x hemisphere (south) | 1.83 | 1.02 | 1.8 | 0.072 |
| Urbanization (suburban) x season (winter) x hemisphere (south) | 1.72 | 1.38 | 1.2 | 0.21 |
| Absolute latitude x urbanization (suburban) x season (winter) x hemisphere (south) | -0.075 | 0.035 | -2.1 | 0.03 |
| Absolute latitude x urbanization (urban) x season (winter) x hemisphere (south) | -0.04 | 0.044 | -0.9 | 0.36 |

**Table S6.** Diversity analyses redone with data compiled at the 5×5 km scale. Response is species richness per 5×5 km grid cell, including only cells with at least 163 eBird checklists. Predictors are as in our main analysis (Table S1). However, given the smaller dataset we dropped interaction terms involving hemisphere to aid model convergence. Model results visualized in Figure S2.

| **Linear model: how urbanization affects year-round bird diversity across latitudes (at 5 km scale)**<br>$\sqrt{\text{species richness}} \sim \text{absolute latitude} \times \text{urbanization} + \text{hemisphere} + \log(\text{number of checklists}) + \text{elevation} + \text{spatial autocovariate}$<br>(df=9 and 1627, $R^2 = 0.66$) | | | | |
| --- | --- | --- | --- | --- |
| Type III ANOVA Results |  |  |  |  |
| Variable | Sum Sq | Df | F value | P value |
| Absolute latitude | 2904.68 | 1 | 1131.66 | < 2E-16 |
| Urbanization | 602.52 | 2 | 117.37 | < 2E-16 |
| Hemisphere | 44.90 | 1 | 17.49 | < 2E-16 |
| Log(number of checklists) | 1381.30 | 1 | 538.15 | < 2E-16 |
| Elevation | 655.35 | 1 | 255.32 | < 2E-16 |
| Spatial autocovariate | 2886.58 | 1 | 1124.61 | < 2E-16 |
| Absolute latitude x urbanization | 278.35 | 2 | 54.22 | < 2E-16 |
| **Linear model: effect of urbanization on bird diversity across latitudes by season (at 5 km scale)**<br>$\sqrt{\text{species richness}} \sim \text{absolute latitude} \times \text{urbanization} \times \text{season} + \text{hemisphere} + \log(\text{number of checklists}) + \text{elevation} + \text{spatial autocovariate}$<br>(df = 22 and 1486 , $R^2 = 0.73$) | | | | |
| Type III ANOVA Results |  |  |  |  |
| Variable | Sum Sq | Df | F value | P value |
| Absolute latitude | 916.67 | 1 | 374.07 | < 2E-16 |
| Urbanization | 292.49 | 2 | 59.68 | < 2E-16 |
| Hemisphere | 207.67 | 1 | 84.74 | 0.002 |
| Season | 22.20 | 1 | 9.06 | < 2E-16 |
| Log(number of checklists) | 933.78 | 1 | 381.05 | < 2E-16 |
| Elevation | 806.79 | 1 | 329.23 | < 2E-16 |
| Spatial autocovariate | 2320.98 | 1 | 947.13 | < 2E-16 |
| Absolute latitude x urbanization | 123.72 | 2 | 25.24 | < 2E-16 |
| Absolute latitude x season | 529.36 | 1 | 216.02 | 1.65E-11 |
| Urbanization x season | 29.44 | 2 | 6.01 | 0.0023 |
| Absolute latitude x urbanization x season | 1.51 | 2 | 0.31 | 0.74 |

**Table S7:**
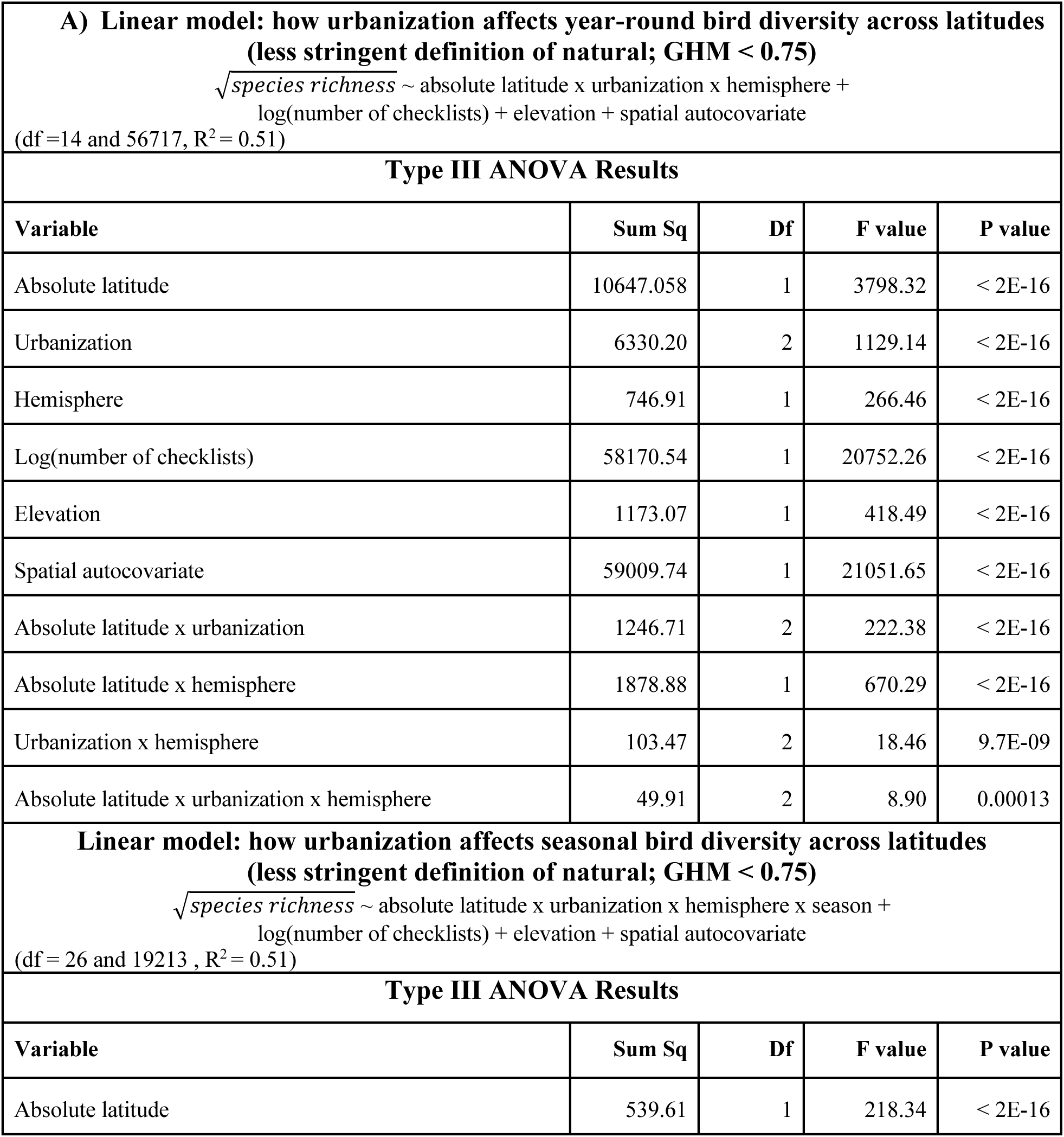
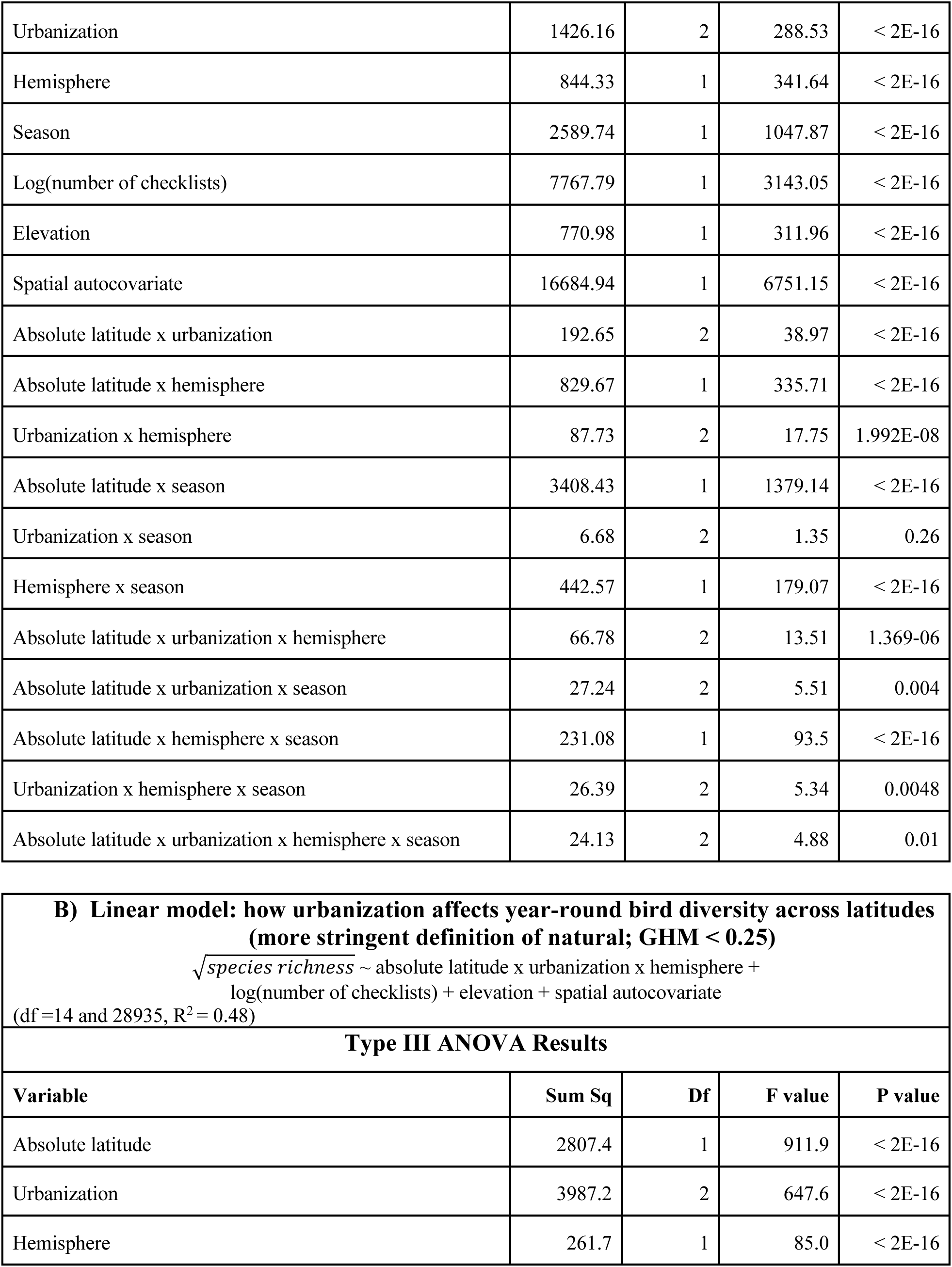
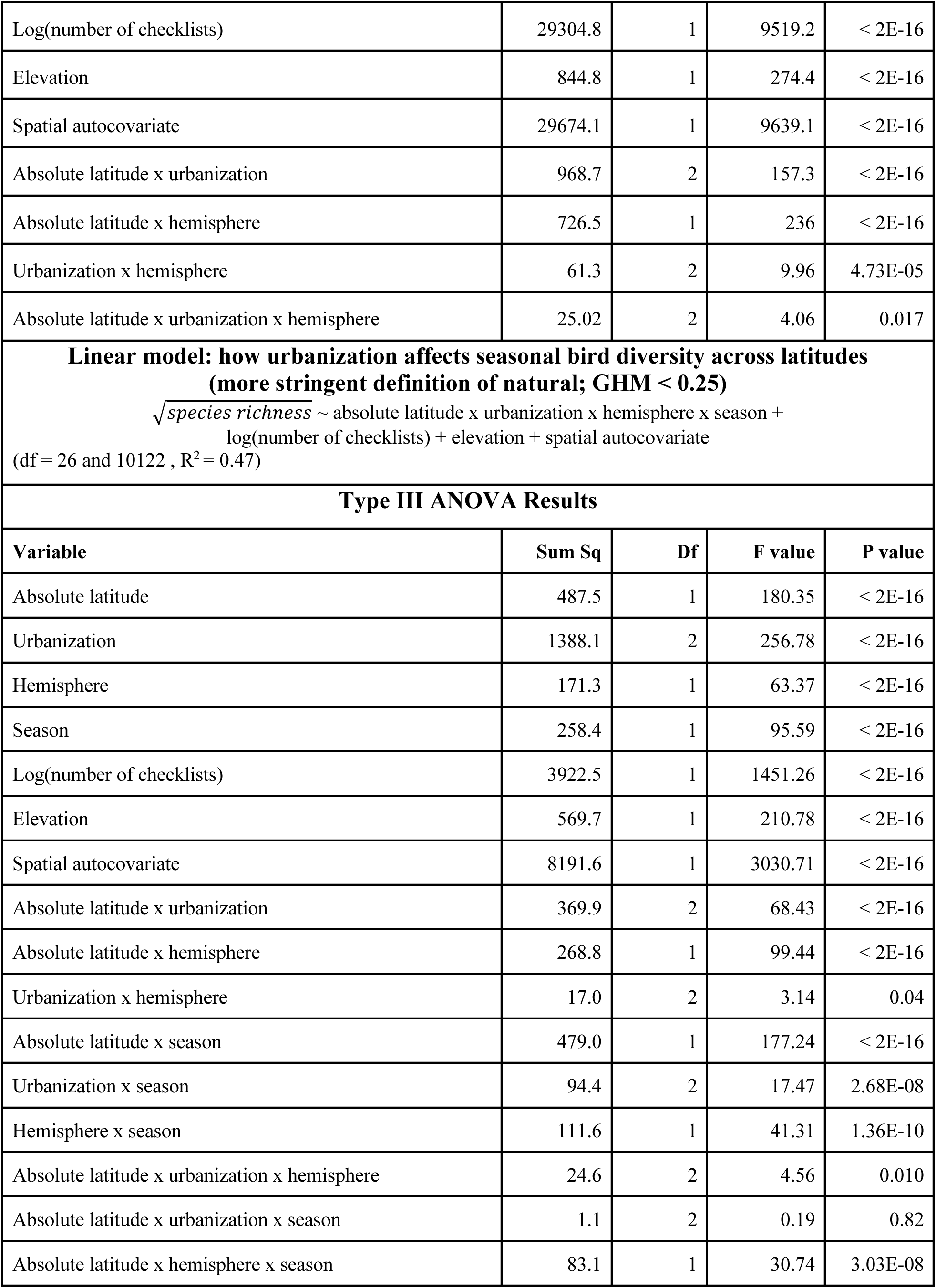
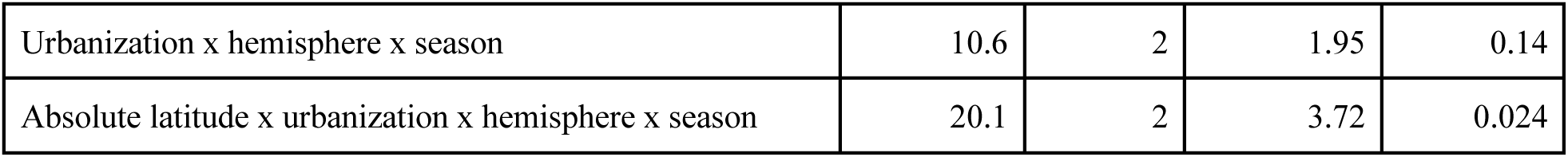
Diversity analyses redone with different thresholds for ‘natural’. Response is species richness per 1×1 km grid cell, as in main diversity analyses in Tables S1 & S5. In main analyses, natural cells are defined as those with <300 human inhabitants and a Global Human Modification (GHM) score of <0.5. A) Analyses redone with a more lenient definition of natural, which includes cells with <300 inhabitants but a GHM score up to <0.75 (i.e. including greater modification). B) Analyses redone with a stricter definition of natural, including only cells with <300 inhabitants and a GHM score <0.25. Predictors are as in Table S1 for year-round analyses and Table S5 for seasonal analyses. Model results are visualized in Figure S3.

**Table S8:**
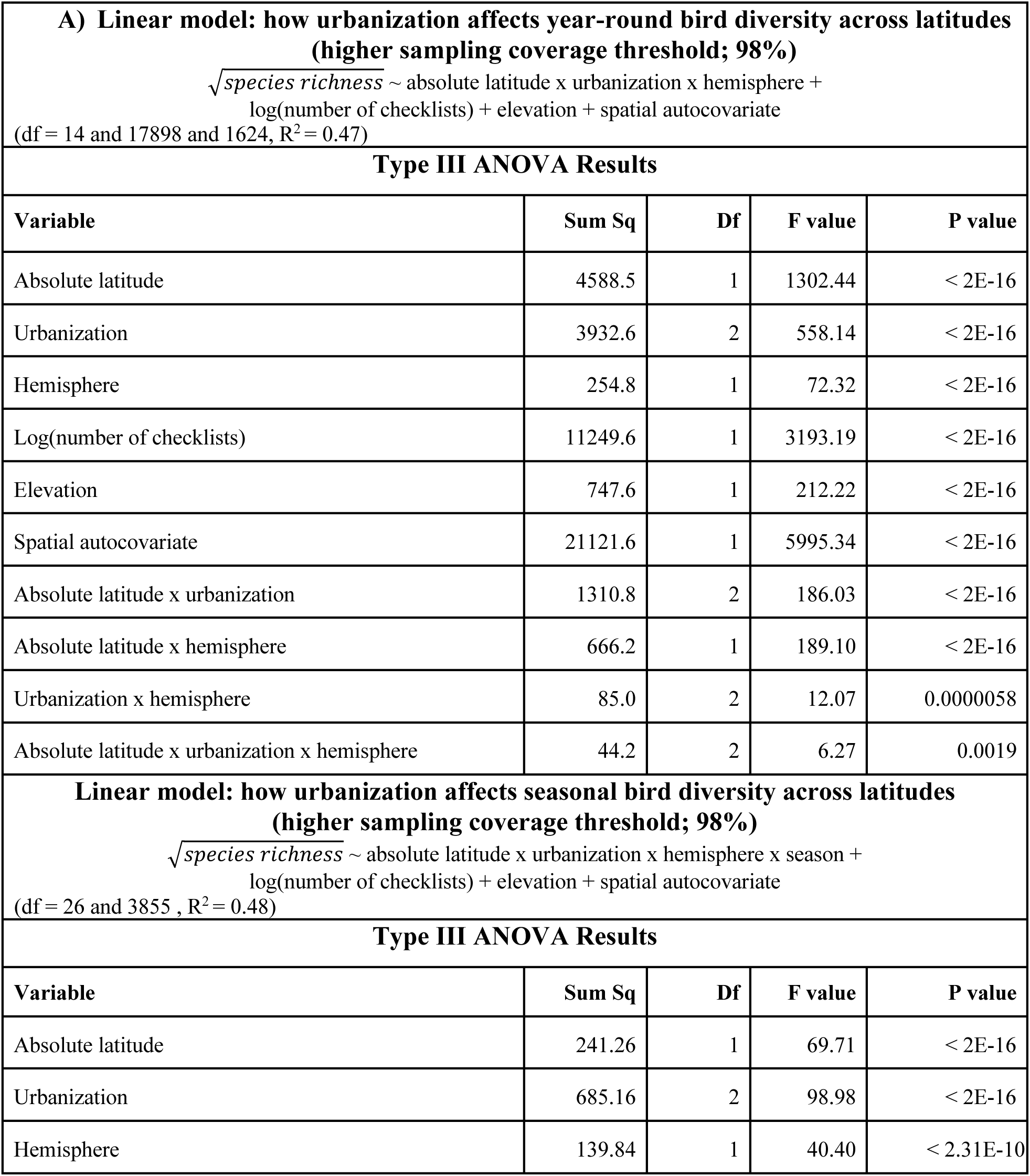
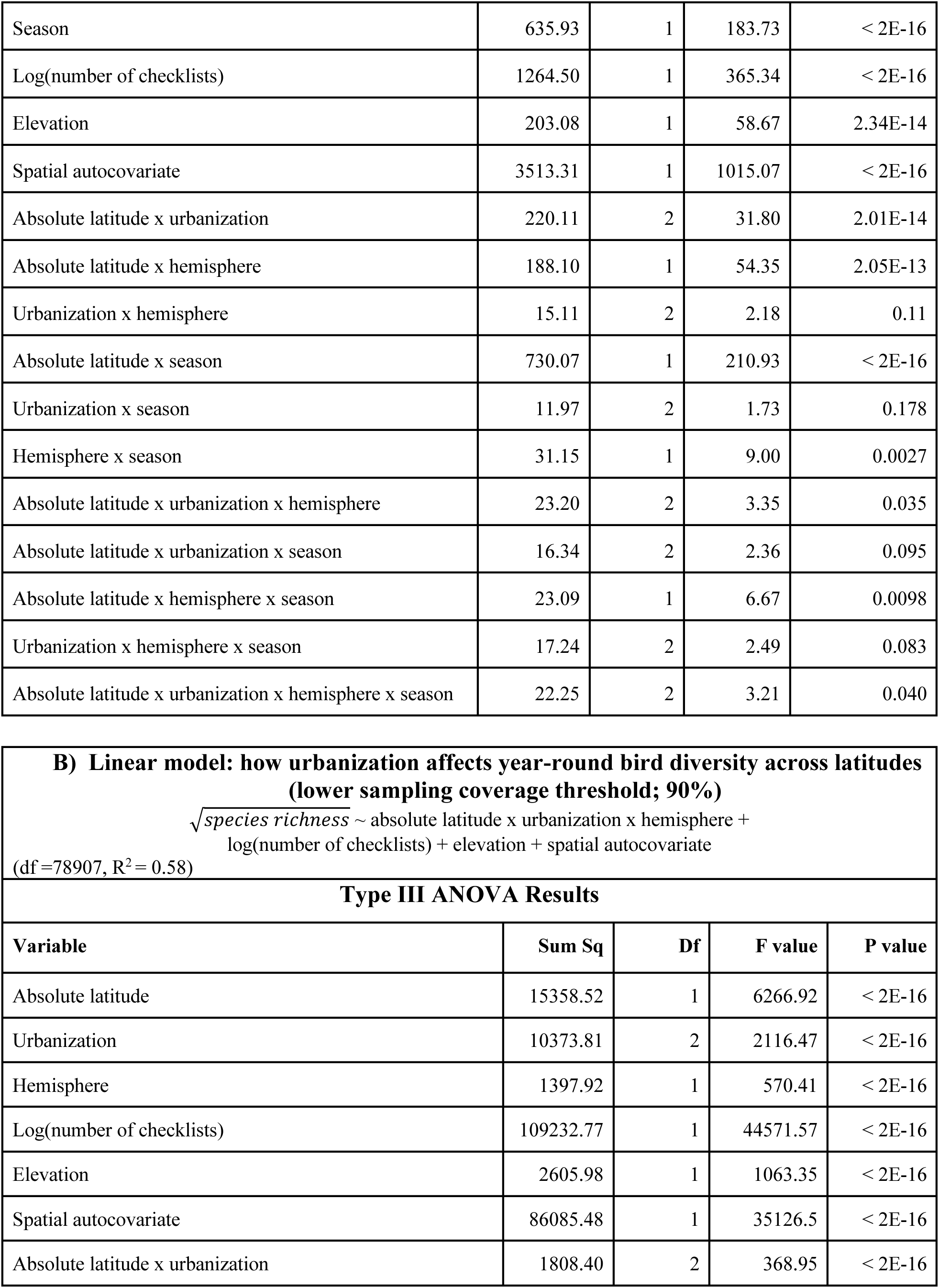
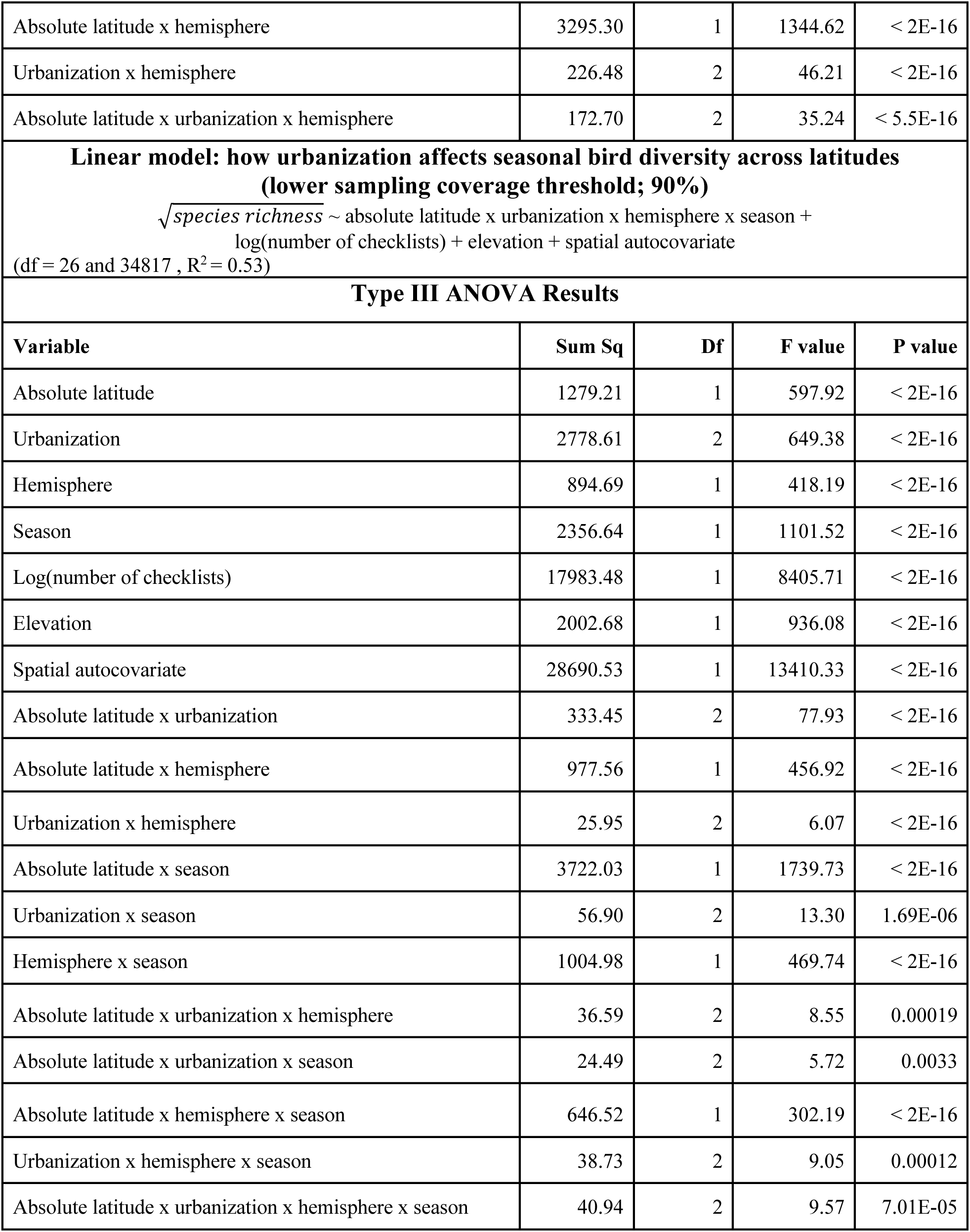
Diversity analyses redone with different thresholds for minimum sampling effort. Minimum sampling intensity in main analyses was determined using the estimated number of eBird checklists needed to reach a 95% sampling coverage in the 500 richest cells; analyses of year-round diversity used the 95th percentile of this distribution, while analyses of seasonal diversity used the 75th percentile. We re-ran analyses using the 95th and 75th percentile of the distribution needed to reach A) 98% sampling coverage (i.e. a stricter threshold), or B) 90% sampling coverage. Models with sample size details are visualized in Figure S4.

**Table S9:** Diversity analyses redone including only natural cells ≤20 km from an urban cell. . We evaluated whether the weaker latitudinal diversity gradient detected in urban areas might be an artefact of urban and natural cells sampling different ecosystems (i.e. if cities are non-randomly built in certain ecosystems and those ecosystems happen to have a naturally weaker latitudinal diversity gradient). To do so, we included only natural cells that were within 20 km of an urban cell, i.e. presumably sampling similar potential bird communities. Model results are visualized in Fig. S7.

| **Linear model: how urbanization affects year-round bird diversity across latitudes (only natural cells within 20 km of an urban cell)**<br>$\sqrt{\text{species richness}} \sim \text{absolute latitude} \times \text{urbanization} \times \text{hemisphere} + \log(\text{number of checklists}) + \text{elevation} + \text{spatial autocovariate}$<br>(df=14 and 26900, $R^2 = 0.45$) | | | | |
| --- | --- | --- | --- | --- |
| Type III ANOVA Results |  |  |  |  |
| Variable | Sum Sq | Df | F value | P value |
| Absolute latitude | 820.63 | 1 | 261.70 | < 2E-16 |
| Urbanization | 1777.49 | 2 | 283.42 | < 2E-16 |
| Hemisphere | 147.63 | 1 | 47.08 | 6.96E-12 |
| Log(number of checklists) | 28796.83 | 1 | 9183.43 | < 2E-16 |
| Elevation | 166.15 | 1 | 52.99 | 3.45E-13 |
| Spatial autocovariate | 21712.57 | 1 | 6924.23 | < 2E-16 |
| Absolute latitude x urbanization | 415.46 | 2 | 66.25 | < 2E-16 |
| Absolute latitude x hemisphere | 314.83 | 1 | 100.40 | < 2E-16 |
| Urbanization x hemisphere | 98.45 | 2 | 15.70 | 0.00000015 |
| Absolute latitude x urbanization x hemisphere | 67.22 | 2 | 10.72 | 0.000022 |
| **Linear model: effect of urbanization on latitudinal diversity across latitudes by season (with natural cells within 20 km of an urban cell)**<br>$\sqrt{\text{species richness}} \sim \text{absolute latitude} \times \text{urbanization} \times \text{hemisphere} \times \text{season} + \log(\text{number of checklists}) + \text{elevation} + \text{spatial autocovariate}$<br>(df= 26 and 8972 , $R^2 = 0.39$) | | | | |
| Type III ANOVA Results |  |  |  |  |
| Variable | Sum Sq | Df | F value | P value |
| Absolute latitude | 33.75 | 1 | 12.18 | 0.00048 |
| Urbanization | 302.5 | 2 | 54.59 | <2E-16 |
| Hemisphere | 43.72 | 1 | 15.78 | 7.16E-05 |
| Season | 70.16 | 1 | 25.32 | 4.95E-07 |
| Log(number of checklists) | 3328.18 | 1 | 1201.19 | < 2E-16 |
| Elevation | 148.97 | 1 | 53.77 | 2.45E-13 |
| Spatial autocovariate | 6235.82 | 1 | 2250.61 | 2E-16 |
| Absolute latitude x urbanization | 47.58 | 2 | 8.59 | 0.00018 |
| Absolute latitude x hemisphere | 53.66 | 1 | 19.37 | 0.052 |
| Urbanization x hemisphere | 16.33 | 2 | 2.95 | 0.052 |
| Absolute latitude x season | 106.6 | 1 | 38.47 | 5.80E-10 |
| Urbanization x season | 34.92 | 2 | 6.3 | 0.0018 |
| Hemisphere x season | 24.06 | 1 | 8.68 | 0.0032 |
| Absolute latitude x urbanization x hemisphere | 12.46 | 2 | 2.25 | 0.105 |
| Absolute latitude x urbanization x season | 8.75 | 2 | 1.58 | 0.21 |
| Absolute latitude x hemisphere x season | 23.21 | 1 | 8.38 | 0.0038 |
| Urbanization x hemisphere x season | 9.94 | 2 | 1.79 | 0.16 |
| Absolute latitude x urbanization x hemisphere x season | 11.75 | 2 | 2.12 | 0.12 |

